# The determinants of genetic diversity in butterflies – Lewontin’s paradox revisited

**DOI:** 10.1101/534123

**Authors:** Alexander Mackintosh, Dominik R. Laetsch, Alexander Hayward, Brian Charlesworth, Martin Waterfall, Roger Vila, Konrad Lohse

## Abstract

Under the neutral theory, genetic diversity is expected to be an increasing function of population size. However, comparative studies have consistently failed to find any strong correlation between measures of census population size and genetic diversity. Instead, a recent comparative study across several animal phyla identified propagule size as the strongest predictor of genetic diversity, suggesting that r-strategists that produce many offspring but invest little in each, have greater long-term effective population sizes. We present a comparison of genome-wide levels of genetic diversity across 38 species of European butterflies (Papilionoidea). We show that across butterflies, genetic diversity varies over an order of magnitude and that this variation cannot be explained by differences in abundance, fecundity, host plant use or geographic range. Instead, we find that genetic diversity is negatively correlated with body size and positively with the length of the genetic map. This suggests that variation in genetic diversity is determined both by fluctuations in *N_e_* and the effect of selection on linked neutral sites.

## Introduction

The genetic diversity segregating within a species is a central quantity; it determines the evolutionary potential of a species, and is, in turn, the outcome of its selective and demographic past. Under the neutral theory [1] genetic diversity is expected to be proportional to the product of the effective population size, *N_e_*, and the per-generation mutation rate, *µ* [2] (provided that *N_e_µ* is sufficiently small that the infinite sites model is applicable [3]). Given that census population size varies widely across the tree of life, much of the variation in genetic diversity between species should be due to differences in census size. In actuality, correlates of census size, such as geographic range, have repeatedly been found to be poor predictors of genetic diversity [4, 5, 6, 7]. In addition, genetic diversity seems to vary remarkably little overall, given the wide range of population sizes seen in nature. While the fact that there are only four alternative states for a nucleotide site suggests a hard upper bound (of 0.75 assuming no mutational bias) on the possible level of nucleotide site diversity [8], neutral genetic diversity in natural populations generally remains well below this. While the extremely narrow ranges of genetic diversity reported by early comparative studies based on allozymes [6] are partly explained by balancing selection [9], diversity at nearly neutral sites is generally restricted to a narrow range of two orders of magnitude [4].

This observation, known as Lewontin’s paradox, has baffled evolutionary biologists for over half a century. Proposed solutions to the paradox are generally of two types; the first proposes that there may be a negative relationship between *N_e_* and *µ* [10], and the second seeks reasons why *N_e_* shows such little variation between species [11]. Given the lack of firm evidence for large differences in mutation rate among species with different levels of variability, recent comparative studies have focused on identifying life-history factors which determine and constrain variation in long term *N_e_* and hence genetic diversity [12]. In particular, Romiguier et al. [5] and Chen et al. [13] have uncovered a striking correlation between reproductive strategy and genetic diversity across the animal kingdom: species that are short-lived and invest little into many offspring (r-strategists) tend to have higher genetic diversity than long-lived species with few offspring and large parental investment (K-strategists). One explanation may be that K-strategists are able to avoid extinction at low population sizes, while r-strategists require much larger populations to buffer against environmental fluctuations.

An alternative (but not mutually exclusive) explanation for the narrow range of genetic diversity observed in nature, is that natural selection continuously removes neutral diversity linked to either beneficial [11] or deleterious variants [14, 15]. Because the efficacy of selection depends on *N_e_s*, selection is expected to be more efficient and therefore remove more neutral linked sites in species with large *N_e_*. Recently, Corbett-Detig et al. [16] have shown that the proportional reduction of neutral diversity due to selection at linked sites does indeed correlate with measures of census size such as geographic range and (negatively) with body size. While Corbett-Detig et al. [16] argue that this can explain”… why neutral diversity does not scale as expected with census size“, a reanalysis of their data [17] concludes that the effect of selection on linked neutral diversity is too small to provide a general explanation for the narrow range of genetic diversity seen in nature. Thus, while recent comparative studies have identified major life history correlates of genetic diversity across the tree of life, and have found support for the idea that selection reduces genetic diversity at linked neutral sites, several questions remain open: What determines variation in genetic diversity across species with similar life-history strategies? Can we identify life history traits other than fecundity that determine a species’ resilience against environmental fluctuations and so correlate with genetic diversity? Does selection merely constrain neutral genetic diversity or can it explain variation?

Here we address these questions using butterflies (Papilionoidea) as a model system. Papilionoidea share a common ancestor approximately 119 million years ago (MYA) [18], and are characterised as r-strategists given their short life span and high fecundity [19]. Butterflies, in particular European species, on which we focus, are arguably the best studied group of insects. Thanks to centuries of study by scientists and amateur naturalists together with numerous recording schemes, butterfly taxonomy, geographic ranges and life-histories are known in great detail. While butterflies exhibit comparatively little variation in life-history strategy, one may still expect fecundity traits (e.g. relative egg size and voltinism, i.e. the number of generations per year) to affect genetic diversity given the strong correlation between fecundity and genetic diversity across animals [5]. Alternatively, if robustness to fluctuations in population size is the ultimate determinant of genetic diversity – as Romiguier et al. argue [5] – one would expect other life history traits to correlate with genetic diversity. In particular, more specialized species may be able to avoid extinction in spite of small census sizes and thus have reduced long-term *N_e_*. While niche breadth is difficult to quantify for many taxa, and therefore has not so far been considered in comparative analyses of genetic diversity, accurate data for the number of larval host plants (LHP) exist for European butterflies.

We estimated genetic diversity from *de novo* transcriptome data for 38 butterfly species (sampling two individuals from each, Supplementary Data 1) and compiled estimates of census size (which we estimated as the product of abundance and geographic range), body size, reproductive output (voltinism and relative egg volume) and the number of LHPs from the literature (Supplementary Data 2, see Methods). Additionally, we tested whether genome size and recombination rate affect genetic diversity. In the absence of detailed recombination maps, we use the number of chromosomes as a proxy for the length of the genetic map. This assumes a map length of 50cM per chromosome and male meiosis on average (given the lack of recombination in females in Lepidoptera), which is supported by linkage maps [20, 21].

To investigate what determines genetic diversity in butterflies, we estimated the relations between seven potential traits (census size, body size, voltinism, relative egg volume, LHP breadth, genome size and chromosome number) and average nucleotide site diversity [22] in a generalized linear mixed model. For simplicity, we restricted the estimation of synonymous diversity to fourfold degenerate sites (*π*_4*D*_), as these genic sites are the least constrained by selection and can be assumed to be nearly neutral. Conversely, non-synonymous diversity was estimated at zero-fold degenerate sites (*π*_0*D*_), i.e. sites where any nucleotide change leads to an amino acid difference. Our rational for modelling *π*_4*D*_ and *π*_0*D*_ jointly was to better understand the nature of the underlying forces at the population level: theory on the effect of selection on neutral diversity predicts that any correlate of neutral genetic diversity (*π*_4*D*_) that increases *N_e_* in the absence of selection should also correlate with non-synonymous diversity (*π*_0*D*_), but do so less strongly [23]. This is because the increase in diversity due to reduced genetic drift is counteracted by the removal of diversity due to more efficient selection. We would therefore expect a weaker correlation for sites that are directly affected by selection than for neutral, linked sites. In contrast, any trait that affects non-synonymous genetic diversity (*π*_0*D*_) via the absolute strength of selection (i.e. affects *s* but not *N_e_*) should be more strongly correlated with diversity at non-synonymous sites (*π*_0*D*_) than synonymous sites (*π*_4*D*_), which are only indirectly affected.

## Results

### Neutral diversity varies over an order of magnitude across butterflies

Genetic diversity was estimated for 38 species of European butterfly from five families: Papilionidae, Hesperiidae, Pieridae, Lycaenidae, Nymphalidae (Fig. 1). For 33 species, we generated and *de novo* assembled short read RNA-seq data for two individuals, for five species raw RNA-seq reads were downloaded from a previous study [5] (Supplementary Data 3). Variants in each species were called by mapping reads back to reference transcriptomes. Only transcripts present in a set of 1314 single-copy orthologues, which we identified from the 33 transcriptomes with high completeness (BUSCO scores 96.3 − 98.4%, Fig. S1), contributed to estimates of genetic diversity.

**Fig 1.**
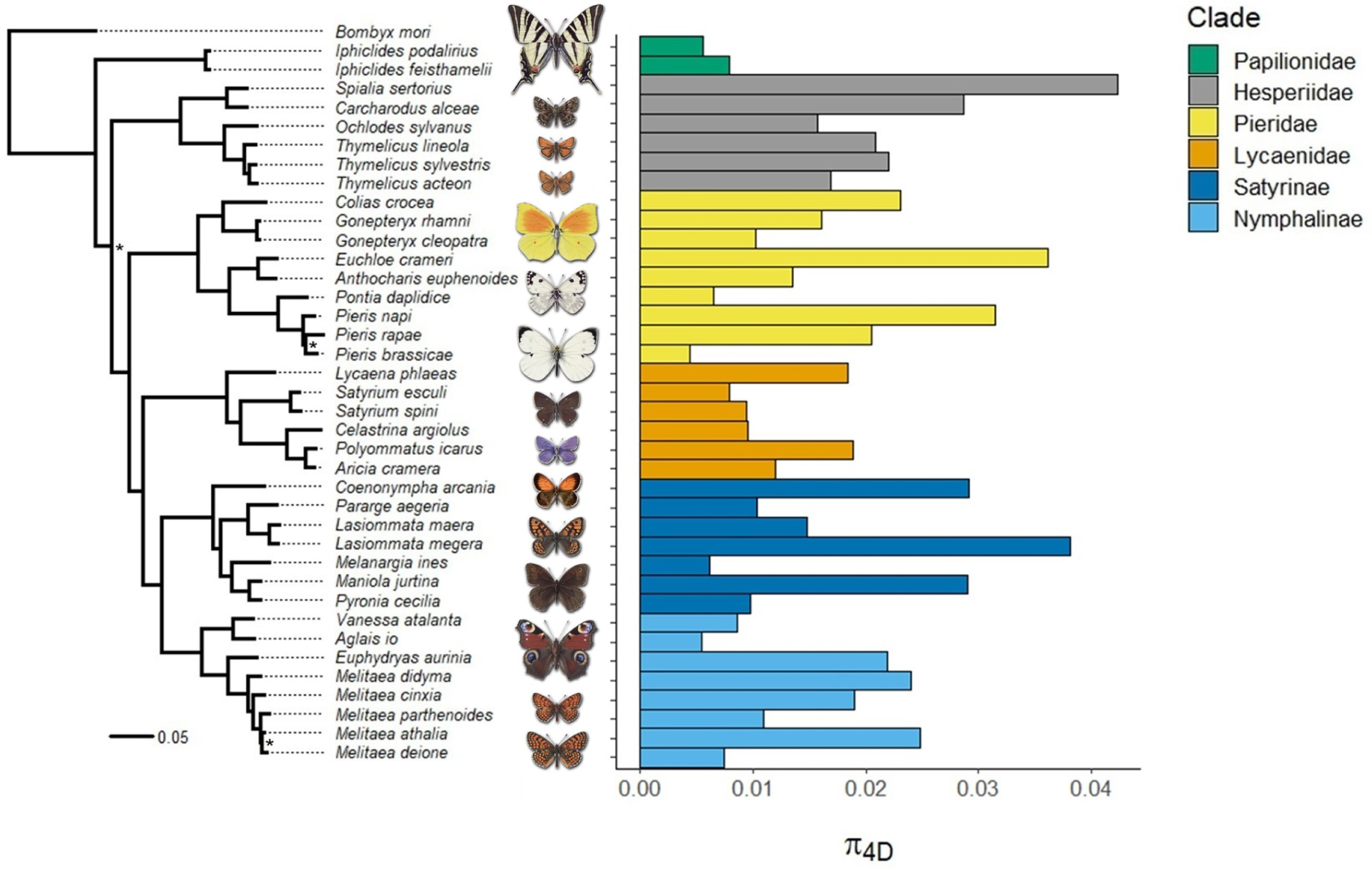
Neutral genetic diversity (*π*_4*D*_) across European butterfly species. The phylogeny is based on 218 single-copy orthologues and rooted with the silkmoth *Bombyx mori* as an outgroup. All nodes have 100% bootstrap support unless marked with an asterisk (70-99%). The barplot on the right shows genome-wide estimates of *π*_4*D*_ for 38 focal species sampled from the six major clades of Papilionoidea present in Europe. The phylogeny explains very little of the variation in *π*_4*D*_ in butterflies.

Mean neutral genetic diversity as measured by *π*_4*D*_ across this set of butterfly species (*π*_4*D*_ = 0.0173) is typical of insects [4, 16, 13]. *π*_4*D*_ varies over an order of magnitude: from 0.0044 in *Pieris brassicae*, the cabbage white, to 0.0423 in *Spialia sertorius*, the red-underwinged skipper (Fig. 1). Assuming neutrality and a per site per generation spontaneous mutation rate of *µ* = 2.9 ×10^−9^ [24], this range corresponds to *N_e_* on the order of 10^5^ to 10^6^ individuals. As expected, this range is much lower than that reported for distantly related animal taxa [5, 13]. While Romiguier et al. [5] - sampling across the entire animal kingdom – found that species in the same taxonomic family have similar genetic diversity, we observed no significant family effect in butterflies (ANOVA, *F*_4,33_ = 2.071, *p* = 0.107). More generally, phylogeny was a poor predictor of neutral genetic diversity in butterflies (*n* = 38, Pagel’s = 5.8 * 10^−5^, *p* = 1.000, assuming that *π*_4*D*_ evolves in a random walk along the phylogeny), Fig. 1).

### Estimates of non-synonymous diversity and the efficacy of selection

Since directional selection will purge (or fix) mutations at non-synonymous sites [25], we expect diversity at these sites to be greatly reduced compared to synonymous sites. Under the nearly neutral theory [26] and assuming a gamma distribution for the distribution of mutational effects on fitness (DFE), the slope of the negative linear relationship between ln(*π*_0*D*_/*π*_4*D*_) and ln(*π*_4*D*_) is equal to the shape parameter, *β* [23]. Within this set of butterfly species,*π*_0*D*_ and *π*_4*D*_ typically differed by an order of magnitude. The slope of the relationship between ln(*π*_0*D*_/*π*_4*D*_) and ln(*π*_4*D*_) (Fig. S2) implies a substantial fraction of weakly deleterious mutations (*β*= 0.44, 95% *CI* = 0.36 − 0.53). This is higher than the estimates for *Heliconius* butterflies (0.08 - 0.28) [13], but compatible with previous estimates of the DFE for *Drosophila* based on the site frequency spectrum [27, 28].

### Nuclear and mitochondrial diversity are uncorrelated

Mitochondrial (mt) genes are an easily accessible source of variation data and have been extensively used to infer the phylogeographic history of species and populations in Europe [29, 30]. However, it is becoming increasingly clear that variation in mt diversity largely reflects selective processes [31] and variation in mt mutation rates rather than rates of genetic drift [32, 33]. In groups with Z/W sex determination, such as butterflies, mt diversity may be additionally reduced by selection acting on the W chromosome (which is co-inherited with the mitochondrion) [34]. Several comparative studies have shown that mt diversity is uncorrelated with measures of abundance and nuclear diversity [35, 32, 33]. We find that across European butterflies mt diversity at the *COI* barcode locus (*π_mt_*) is only very weakly and non-significantly correlated with genome-wide neutral diversity *π*_4*D*_ (Pearson’s correlation, *d.f.* = 36 *r* = 0.1523, *p* = 0.362) and *π*_0*D*_ (Pearson’s correlation, *d.f.* = 36 *r* = 0.268, *p* = 0.104) (Fig. S3).

### Census population size, host plant breadth and reproductive strategy are uncorrelated with genetic diversity

We find that census population size is uncorrelated with both *π*_0*D*_ and *π*_4*D*_ (Table S1). This suggests that present day ranges and abundance have little to do with long term *N_e_* in butterflies. Unlike recent studies which have found that propagule size strongly correlates with neutral genetic diversity across much wider taxonomic scales [5, 13], we find no significant effect of relative egg size (egg volume / body size) on *π*_4*D*_ (Table S1). Similarly, voltinism is uncorrelated with *π*_4*D*_ (*p* = 0.151, Table S1). Although not significant, the trend of polyvoltine taxa having greater *π*_4*D*_ is at least consistent with the idea that r-strategists have larger long-term *N_e_* [5]. We find that larval hosts plant (LHP) breadth has no significant effect on *π*_4*D*_ or *π*_0*D*_ (Table S1). This is true regardless of whether we classify species as monophagous if all LHPs are within one family (and polyphagous otherwise) or instead consider the number of LHP species as a predictor (Fig. S4).

Only one trait, body size, is significantly and negatively correlated with *π*_4*D*_ (*p* < 0.005, Table 1, Fig. 2A): smaller butterfly species tend to have higher genetic diversity. As predicted for correlates of long term *N_e_*, the effect is weaker for *π*_0*D*_ (Table 1) than *π*_4*D*_. We can express the effects of body size on ln(*π*_4*D*_) and ln(*π*_0*D*_) in terms of ln(*π*_0*D*_/*π*_4*D*_). This ratio is weakly and positively correlated with body size (posterior mean slope = 0.123, *p* = 0.049), suggesting that selection is more efficient in smaller species.

**Table 1.** Posterior mean estimates of the slope of linear correlates of genetic diversity at synonymous (*π*_4*D*_) and non-synonymous (*π*_0*D*_) sites inferred under a minimal model.

| Predictor | Response | Posterior mean slope | 95% CI | $p_{MCMC}$ |
| --- | --- | --- | --- | --- |
| Body size | $\pi_{4D}$ | -0.321 | -0.513, -0.110 | 0.002 |
| Body size | $\pi_{0D}$ | -0.200 | -0.335, -0.064 | 0.005 |
| Chrom. number | $\pi_{4D}$ | 0.277 | 0.103, 0.468 | 0.003 |
| Chrom. number | $\pi_{0D}$ | 0.150 | 0.024, 0.267 | 0.017 |

**Fig 2.**
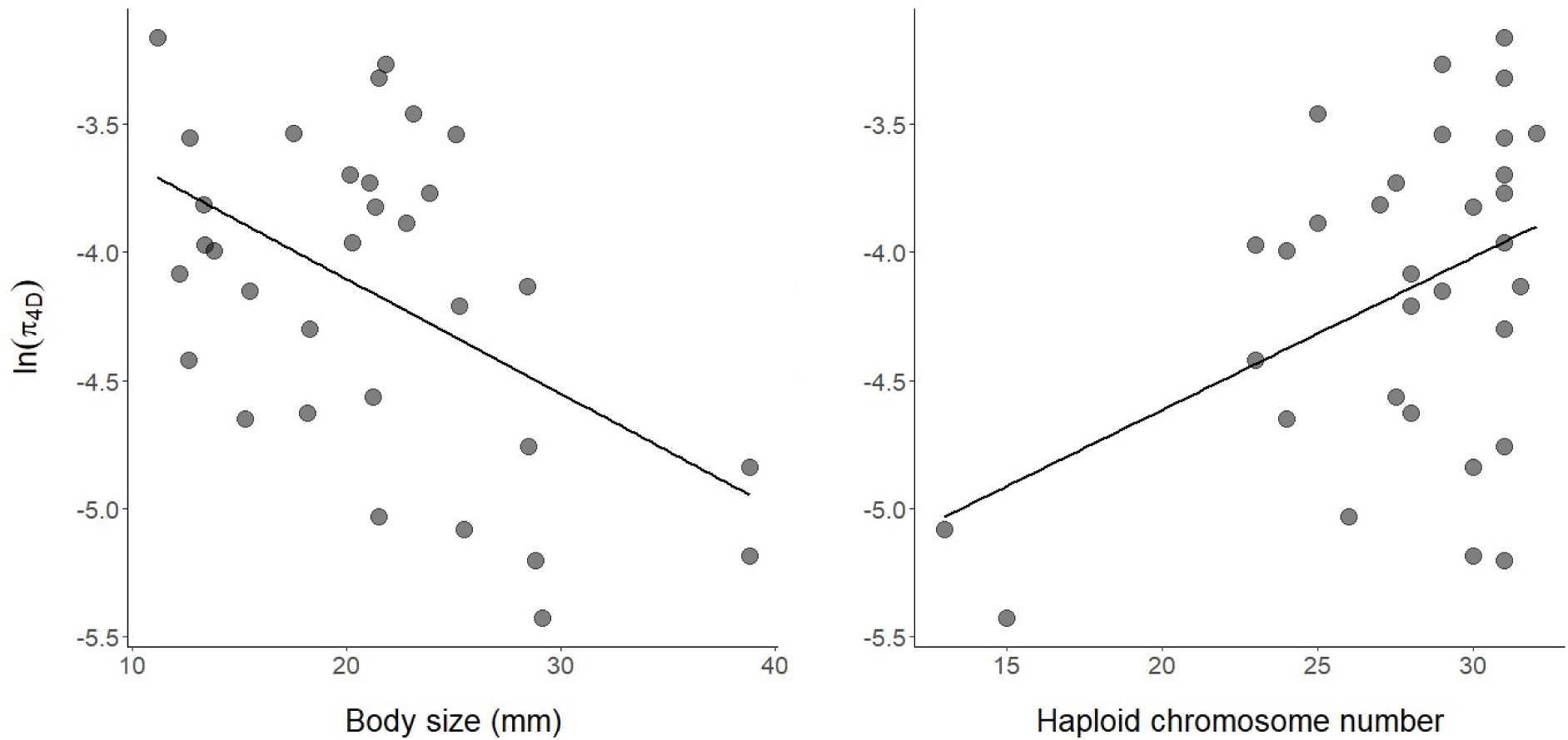
Neutral diversity *π*_4*D*_ in butterflies is negatively correlated with body size (left) and positively with the number of chromosomes (right).

### Chromosome number, but not genome size, correlates with genetic diversity

While *ln*(*π*_4*D*_) correlates positively and significantly with chromosome number (posterior mean slope = 0.277, *p* = 0.003, Table 1, Fig. 2B), it is uncorrelated with genome size, i.e. the physical length of the genome (estimated using flow cytometry, see Methods) (Table S1). Assuming that the number of genes in the genome (and other potential targets of selection) is more or less constant and independent of genome size, population genetic theory predicts the aggregate effect of selection on linked neutral diversity to be largely determined by the map length of a chromosome, for a given set of selection and mutation parameters [11, 15, 14, 36] (see Discussion).

Although (unsurprisingly) the effect of chromosome number depends disproportionately on the two species with the fewest chromosomes (*Pieris brassicae*, *n_c_* = 15, and *Melanargia ines*, *n_c_* = 13, Fig. 2B), removing both species gives a very similar estimate of the slope (posterior mean slope = 0.284, *p* = 0.128).

### Pleistocene bottlenecks and demography

Genetic diversity in many European taxa has been shaped by the cycles of isolation into, and range expansion out of, glacial refugia during the Pleistocene [29, 30, 37]. While we have sought to minimize the effects of Pleistocene history by focusing sampling on a single Pleistocene refugium, Iberia, our inferences could be confounded in at least two ways: Firstly, rather than being solely driven by long term *N_e_*, variation in genetic diversity in Iberia may be affected by gene flow from other refugia [38] or even species [39]. Secondly, even if Iberian populations are little affected by admixture, they may have undergone drastic (and potentially different) changes in *N_e_* in response to past climatic events. Population bottlenecks affect *π*, but correspond to a sudden burst in coalescence rather than a change in the long term rate [40]. Population bottlenecks would also affect our interpretation of *π*_0*D*_/*π*_4*D*_ as a measure of the efficacy of selection. Since *π*_0*D*_ recovers more quickly than *π*_4*D*_ after a bottleneck, one would expect taxa that have undergone recent changes in *N_e_* to fall above the line in *ln*(*π*_4*D*_) vs *ln*(*π*_0_*/*π**_4*D*_) correlation (Fig. S2).

While modelling demography from transcriptome data is challenging, the distribution of heterozygous sites in a single diploid individual contains some information about past demography. In particular, an extreme bottleneck or a rapid expanding lead to strongly correlated pairwise coalescence times. Considering a fixed length of sequence, we expect the number of heterozygous sites *S* to be Poisson distributed, whereas intermediate bottlenecks result in multimodal distribution of *S* with an increased variance relative to a constant sized population [41]. However, the majority of species show a unimodal, long tailed distribution of *S*, more akin to that expected for a population of constant *N_e_* than the limiting case of an extremely bottlenecked (or rapidly expanding) population. In fact, only seven species have a higher variance in *S* than expected for population of constant size (Fig. S6 and S7).

### Robustness to population structure

The relationship between genetic diversity and population size predicted by the neutral theory assumes a randomly mating population at mutation-drift equilibrium. Since population structure is ubiquitous, an obvious question is to what extent our findings are confounded by differences in population structure across species. For example, the correlation between body size and diversity may simply be a consequence of the reduced dispersal ability in smaller species. If this were the case, we would expect genetic differentiation to also correlate with body size. However, we find no evidence for this: differentiation between individuals sampled *>*500 km apart is low overall (median *F_IT_* = 0.019) and uncorrelated with body size (Pearson’s correlation, *p* = 0.804) (Fig. S5). Furthermore, the effect of body size on genetic diversity remains essentially unchanged if we estimate *π*_4*D*_ and *π*_0*D*_ within rather than between individuals. Increased population structure in smaller species can therefore not explain the negative relationship between genetic diversity and body size.

Our dataset does include a handful of species with notably high *F_IT_* within Iberia, such as the Marsh Fritillary *Euphydryas aurinia* and the Pearly Heath *Coenonympha arcania* (*F_IT_* = 0.281 and 0.122, respectively). Interestingly, both species fall above the line of best fit in Fig. S2 suggesting that selection is less efficient globally (i.e. ln(*π*_0*D*_/*π*_4*D*_) is higher) in these species. The presence of different locally adapted subspecies or populations could further increase ln(*π*_0*D*_/*π*_4*D*_). For *E. aurinia* at least three ecotypes/subspecies exist in the Iberian peninsula, but their exact distribution and significance is uncertain. In contrast, the migratory species *Vanessa atalanta* is an outlier in the opposite direction and has lower diversity at non-synonymous sites (*π*_0*D*_) than expected given its neutral diversity (*π*_4*D*_) (Fig. S2).

## Discussion

We have shown that neutral genetic diversity in European butterflies varies over an order of magnitude and that this variation is uncorrelated with both current abundance and several key life-history traits. In particular, and in contrast to previous comparative studies across larger taxonomic scales [5, 13], we do not find any relationship between propagule size or longevity and genetic diversity. We also find little support for the idea that generalist species have larger long term *N_e_* and hence greater levels of genetic diversity. Instead, body size and chromosome number were the only significant correlates of neutral genetic diversity and, together, explain 45% of the variation in genetic diversity across European butterflies. The negative correlation between body size and genetic diversity is consistent with body size limiting population density [42] and therefore long-term *N_e_*. This relationship is not exclusive to butterflies, and has been found in mammals [43] and across animals [5] more widely.

As we show below, the positive correlation between chromosome number and neutral genetic diversity is an expected consequence of selection and mirrors the nearly ubiquitous intraspecific correlation between genetic diversity and recombination rate [44, 45]. Thus, unlike previous comparative studies which have shown that selection merely constrains variation in genetic diversity [16], our results demonstrate that the effect of selection on linked neutral diversity may explain variation in genetic diversity between taxa that differ in the length of the genetic map.

### Niche breadth

The lack of any correlation between estimates of census size and *π*_4*D*_ we find mirrors results of previous studies [5, 4, 7, 6] and suggests that current abundance does not reflect long term *N_e_* in European butterflies. While the distribution of heterozygosity suggests that it is unlikely that variation in genetic diversity across butterflies is due to drastic demographic events during the Pleistocene, very recent demographic changes could explain the weak relationship between estimates of census population size and *π*_4*D*_. In particular, the low genetic diversity of *Pieris brassicae*, a pest species with enormous current population sizes, is compatible with a rapid expansion which may have happened too recently to leave much signal in the data: *V ar*[*S*] is not particularly low for *P. brassicae* (5.67 compared to the mean among species of 4.85). Interestingly, analysis of RAD-seq data from the closely related species *P. rapae* suggests a population expansion ≈ 20,000 yBP (shortly after the last glacial maximum) followed by divergence into subspecies 1200 yBP when *Brassica* cultivation intensified [46]. It is therefore possible that, by contrast, the ancestral *P. brassicae* population remained small after the glacial maximum and only expanded as recently as ≈ 1200 yBP.

If variation in carrying capacity shapes genetic diversity in butterflies, it is perhaps surprising that niche breadth, the number of larval host plants (LHPs), is uncorrelated with *π*_4*D*_. However, given that LHPs vary drastically in geographic range and density, the number of LHPs may be a very crude predictor of a species long-term census size: a species with a single LHP may have very large populations if its host is widespread. Conversely, a generalist may have low long-term *N_e_* due to other biotic factors. For example, *C. argiolus* one of the most widespread and generalist (*>* 100 LHPs) species in our set has relatively low neutral genetic diversity (*π*_4*D*_ = 0.0095).

There are several potential life history traits that might have large effects on long term *N_e_* which we have not considered: in particular, how (in what life-cycle stage) and where species hibernate, the rate of parasitoid attack and the degree of migratory versus sedentary behaviour. Exploring whether these correlate with genetic diversity will require larger sets of taxa.

### Can selection explain the correlation between genetic diversity and chromosome number?

Body size and chromosome number together explain 45% of the variation in genetic diversity across European butterflies (Fig. 2). We have assumed linear relationships between these predictors and genetic diversity without paying any attention to the causative forces at the population level. To gain some insight into whether the genome wide effects of background selection (BGS) [14, 15] and recurrent selective sweeps [11, 47, 36] can plausibly explain the observed relationship between diversity and chromosome number, it is helpful to consider analytic results for the reduction in neutral diversity caused by selection at sites linked to a focal neutral site. We take as a starting point the expression of Coop (2016) [17, eq. 1] for the expected genetic diversity given BGS and sweeps occurring homogeneously along the genome. Note that this approximate result assumes independence between selective events and is based on a considerable body of previous population genetics theory [47, 36, 48] (see Supplementary Data S4 for details):

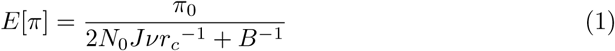

where *π*_0_ = 4*N*_0_*µ* is the genetic diversity in the absence of selection, *B* is the effect of BGS, *v* and *r_c_* are the rates of sweeps and recombination per base pair respectively, in the genomic region under consideration. *J* captures the probability of sweeps leading to coalescence at linked neutral sites. Assuming semi-dominance with selection coefficient *s* in homozygotes for a beneficial mutation, *J ≈ s/*[2 ln (2*N_e_s*)] (for details see Supplementary Data 4). We can think of 2*N*_0_*Jvr_c_^−1^*^1^ as the rate of sweep-induced pairwise coalescence events relative to genetic drift, i.e. in units of 2*N*_0_ generations.

A simple approximation for the effect of BGS is *B*(*r*)≈ exp(*U/r_c_*) [15], where *U* is the per base rate of deleterious mutations per diploid genome. Thus both the effects of BGS and positive selection depend on the ratio of mutational input to recombination rate. We can scale the rates of deleterious mutations and selective sweeps per genome (rather than per bp): assuming that the number of selective targets is fixed across species, the terms for BGS and positive selection are functions of the genetic map length (this assumes a linear relationship between recombination rate and map length), i.e. the number of chromosomes, *n_c_*: *v/r* ≈ 4*v_T_/n_c_* and *B* ≈ exp(*π*4*U_T_/n_c_*), where *v_T_* and *U_T_* are the total numbers of selective sweeps per unit coalescence time and new deleterious mutations per genome respectively. Note that we assume on average half a crossover event per chromosome and male meiosis given the absence of recombination in female Lepidoptera so that the total map length of the genome is 0.25 × *n_c_*.

One immediate conclusion from the above is that, given the large number of chromosomes in butterflies (13 ≤ *n_c_* ≤ 31), BGS can only have a modest effects on neutral diversity: even if we assume a rate of *U_T_* = 1 deleterious mutation per genome, the reduction in diversity due to BGS, *B*(*r*), only ranges between 0.73 and 0.88 for our dataset. Ignoring the effect of BGS, we have:

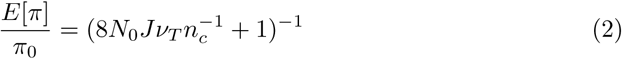

We can use eq. 2 to ask how compatible the expected effect of selective sweeps on neutral diversity is with the estimate of the slope of the relation between ln(*π*_4*D*_) and *n_c_* (Table 1). In the limit of high *v_T_*, eq. 2 implies that 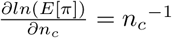, so assuming an average of *n_c_* = 25 chromosomes, we would expect a maximum slope of 0.04. This expected effect of positive selection is compatible with our empirical estimates of the slope between ln(*π*_4*D*_) and *n_c_* (the estimate in Table 1 corresponds to 0.0620 (95 % CI 0.0224, 0.01041) on the untransformed *n_c_*) but requires an extremely high rate of sweeps.

In principle, one can go one step further and use eq. 2 to estimate the rate of sweeps from the data by minimizing the sum of squared differences between observed and predicted *π*_4*D*_ across species. For example, if we assume that *N*_0_ depends linearly on body size, a spontaneous mutation rate of *µ* = 2.9 × 10^−9^ [24], and *J* = 10^−5^ (which corresponds to *N*_0_ = 10^6^ and *s* = 10^−4^ which is consistent with values estimated in insects [27]), we can co-estimate both the correlation between *N*_0_ and body size and *v_T_* (Supplementary Data 5). The best fitting selection regime implies an extremely high rate of sweeps of (*v_T_* ≈ 0.135 per generation). However, this approximate model of the effect of selective sweeps on *π* only fits marginally better than the linear model assumed by `MCMCglmm` (the sum of least squares between the observed set of *π*_4*D*_ estimates and the expected values is 0.00222 and *S* = 0.00226 respectively) and predicts a much narrower range of *π*_4*D*_ than is observed (Fig. S8). Thus, the above calculation agrees with the analysis of Coop [17], in that it shows that simple approximations for the effect of selection on neutral diversity provide a poor fit to the observed overall variation among species in genetic diversity.

### The evolution of chromosome number and genome size

We have so far assumed that chromosome number is simply a proxy for the genetic map length and affects genetic diversity by modulating the effect of selection on linked neutral sites. However, what is cause and effect is far from clear and chromosome number may itself depend on the efficacy of selection. Recently Hill et al. [49] found that chromosomes in *Pieris napi* are derived from multiple ancestral syntenic blocks, suggesting a series of fission events that was followed by the creation of a novel chromosome organisation through fusions. Because *P. napi* returned to a karyotype close to the ancestral *n_c_* = 31 of butterflies, there appears to be some selective advantage in organising the genome this way, and chromosome rearrangements that produce karyotypes distant from *n_c_* = 31 may only be tolerated in populations dominated by drift. There is evidence that chromosomal fusions accumulate in small populations of mammals [50, 51] and in selfing plants [52]. Thus, an alternative explanation for the positive correlation between chromosome number and genetic diversity we find could be that species with low *N_e_* accumulated mildly deleterious chromosome rearrangements through drift. *Pieris brassicae* (*n_c_* = 15) and *Melanargia ines* (*n_c_* = 13) having most likely undergone relatively recent chromosomal fusions (given that in both cases relatives in the same genus have higher *n_c_*) would be consistent with this. As no species within our dataset has *n_c_* >> 31, we cannot test whether the relationship between genetic diversity and chromosome number is quadratic, and thus consistent with a model where reduced *N_e_* may lead to both increases or decreases in *n_c_*. Interestingly, species in the genus *Leptidea*, which have undergone a drastic and recent explosion in chromosome number (*n_c_* ranges between 26 and 120 [53]), appear to have very low genome-wide diversity (*π* across all site between 0.0011 and 0.0038) [54] which is consistent with the idea that extreme karyotypes arise during periods of low *N_e_*.

Lynch & Conery [55] have put forward analogous arguments for the evolution of genome sizes, arguing that genomes may expand in populations with low *N_e_*, if selection against transposable element proliferation and intron expansion becomes inefficient. We find no support for any relationship between genome size and neutral diversity in butterflies. Instead our analyses clearly show that genome size has significant phylogenetic signal across butterflies (*n* = 37, Pagel’s *λ* = 1.000, *p* = 6.1 ∗ 10^−7^) and so must evolve slowly, whereas variation in genetic diversity has little phylogenetic structure (Fig. 1).

### Outlook

While we have only considered a small number of life-history traits and genomic parameters, and have modelled neither the effects of selection nor demography explicitly, it is encouraging that we have identified two simple determinants, which together explain a substantial fraction of the variance in genetic diversity across butterflies. It is clear however, that a more complete understanding of the processes that shape genetic diversity and how these correlate with life-history will require modelling both the demographic and the selective past [56] explicitly. For example, a previous comparative study based on whole genome data reconstructed the directional histories of divergence and admixture between refugial populations for a different guild of insects [38] and found a trend of refugial population being younger in specialist species. An important next step is to include models of selection and its effects on linked sequence in such inferences. Given sufficiently large samples of taxa, one can then tease apart life history traits that affect genetic diversity via demographic parameters (*N_e_* in the absence of selection, gene flow between populations) from those that determine the strength of selection itself. Rather than focusing on pairwise *π*, the most drastic summary of genetic variation, such inferences will require methods that make use of the detailed information contained in genomic data. Another important source of information, which has been exploited by Corbett-Detig et al. [16], but is currently unavailable for most taxa, is provided by direct estimates of the recombination map. Given the detailed knowledge of their taxonomy, ecology, geographic range and their relatively compact genomes, butterflies are perhaps the best test case for attempting a reconstruction of the evolutionary processes that would result in Lewontin’s paradox.

## Methods and Materials

### Sampling and sequencing

Butterflies were hand-netted at various locations across four regions in Iberia (Southern Portugal, Northern Portugal, Catalonia and Asturias, Supplementary Data 1), frozen alive in a liquid nitrogen dry shipper and stored at −80 °C. Two individuals per species were selected for RNA extraction and sequencing. Each species was represented by one female and one male individual when possible. Species identities were confirmed by amplifying and sequencing the standard mitochondrial barcode (a 658-bp fragment of *COI*, primers LepF and LepR [37]) and comparison against a reference database for Iberian butterflies [37] in the following species: *Carcharodus alcae*, *Coenonympha arcania*, *Euphydryas aurinia*, *Melitaea deione Thymelicus acteon* and *T. sylvestris*.

RNA was extracted using a TRIzol (Ambion) protocol according to the manufacturer’s instructions. TruSeq stranded polyA-selected RNA libraries were prepared by Edinburgh Genomics and strand specific 75b paired-end reads were generated on a HiSeq4000 Illumina instrument. Raw reads are deposited at the European Nucleotide Archive (PRJEB31360). RNA-seq datasets for *Melitaea athalia*, *M. cinxia*, *M. didyma*, *M. parthenoides*, and *Thymelicus lineola* — previously analysed in [5] — were retrieved from the European Nucleotide Archive (ENA).

### Data QC and *de novo* transcriptome assembly

Raw read quality was evaluated using `FastQC v0.11.7` [57] and visualised with `MultiQC v1.5a` [58]. Illumina adapters were removed and reads trimmed using `Trimmomatic` [59] (under default parameters) and only reads of length ≥25b were retained. Transcriptomes were assembled *de novo* from both individuals of each species with `Trinity` [60] and are deposited at the XXX database. Assembly completeness was assessed using `BUSCO v3.0.2` [61] together with the Insecta database `insectodb9` (1658 single copy orthologues from 42 insect species) as a reference (Fig. S1, Supplementary Data 2).

### Variant calling

Protein coding transcripts were identified using `Transdecoder` [62], `BLAST` [63] and H`MMER` [64]. Transdecoder was used to find open reading frames (ORFs) within transcripts, while homology searches were done using `BLAST` and `HMMER` to identify transcripts containing known sequences and domains. Finally, the predict function in `Transdecoder` was used to score likely protein coding transcripts based on ORF presence and homology. For each species, reads of both individuals were separately mapped to the longest isoforms of complete proteins using `BWA MEM` [65]. Loci which were suitable for variant calling were selected using the callable loci function in `GATK` [66]. We selected sites with a read depth ≥ 10 and MQ ≥ 1. Callable loci were intersected between individuals using BEDTools [67], removing sites that were not expressed by both individuals in each species. Variants were called using `Freebayes` [68], and only retained if supported by more than three reads, with the variant being found in both the 3’ and 5’ end of reads, as well as in both forward and reverse reads. Excluded variants were masked for downstream analysis, and so did not contribute to the total length or variant count.

### Estimating genetic diversity

Protein clustering using `Orthofinder` [69] revealed 1314 single copy orthologue clusters in the 33 transcriptomes with high completeness (`BUSCO` scores 96.3 - 98.4%). Only the transcripts corresponding to these proteins were used to estimate *π* in each species transcriptome. To minimize the confounding effect of population structure (and inbreeding) we calculated *π_b_*, i.e. the genetic diversity between the two individuals sampled for each species (analogous to *d_XY_*):

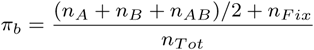

where *n_A_*, *n_B_* are the numbers heterozygous sites unique to individual A and B, and *n_AB_* is the count of shared heterozygous sites and *n_fix_* is the number of fixed differences. Calculations were carried out separately for four-fold degenerate (*π*_4*D*_) and zero-fold degenerate (*π*_0*D*_) sites using the script `bob.py` deposited at www.github.com/DRL/mackintosh2019.

Mitochondrial *π* was calculated for the COI barcode locus using sequences retrieved from the BOLD systems database [70]. Alignments of 658bp for each species were produced in `Bioedit` [71] using CLUSTAL-W [72] and manual inspection. Mean pairwise *π* of each alignment was calculated in `MEGA7` [73].

### Phylogeny reconstruction

Single-copy orthologous proteins present in all transcriptome assemblies — as well as the genome of the silkmoth *Bombyx mori* — were identified with `Orthofinder`. The resulting 218 protein sequences were concatenated for each species, aligned using `MAFFT` [74], and trimmed using `trimAl` [75]. The final alignment contained 59747 amino acid sites, 22429 of which were informative for phylogenetic inference. 20 maximum likelihood (ML) tree searches were conducted using the substitution model PROTGTR+GAMMA, with `RAxML` [76]. To assess confidence in the ML tree, non-parametric bootstrap values were obtained from 100 replicates.

### Statistical analysis

Phylogenetic mixed models were constructed using the R package `MCMCglmm` [77]. Models were bivariate, that is, included two responses, ln(*π*_4*D*_) and ln(*π*_0*D*_), which were assumed to covary and follow a Gaussian distribution. Only the 32 species with data for all seven predictors were included.

Fixed effects were z-transformed when continuous so that estimated effect sizes were comparable for a given response. Phylogeny was included in the model as a random effect based on the inverse matrix of branch lengths in the maximum likelihood species phylogeny (Fig. 1). We assumed the following parameter expanded priors for the random effect (G) and the residual variance (R):

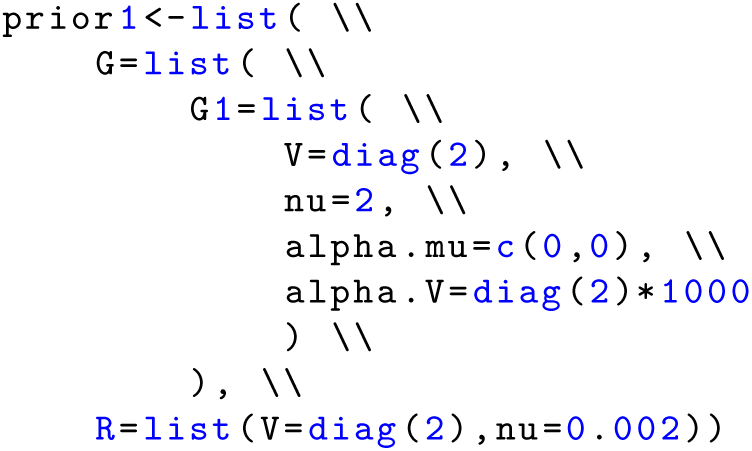

, where *V* is the variance matrix, *ν* the degree of belief, *αV*the covariance matrix, and *αµ* the prior mean. A maximal model, containing all seven predictors as fixed effects, was constructed and then simplified by backwards elimination of predictors. The minimal model therefore only contains predictors with a significant (*α* ≤ 0.05) effect.

### Estimating genome size by flow cytometry

To estimate the size of the genome for each species we followed the protocol outlined by [78], with some minor modifications. In short, head tissue of butterflies (frozen fresh and preserved at −80 °C) were ground in Gailbraith’s buffer and filtered through a 40 µm mesh, resulting in a suspension of free nuclei with minimal cell debris. The solution was centrifuged at 350/500g for 1 minute, then the pellet of nuclei was resuspended in 300*µ*l propidium idodide (50*µ*g/ml; Sigma-Aldrich) and RNAse A (100*µ*g/ml; Sigma-Alrich) for staining and removal of RNA. After 1-2 hours, fluorescence was measured using a BD LSR Fortessa running `Diva v8.0.1`. DNA content of cells were evaluated by propidium iodide binding using a 561nm excitation laser and fluorescence emission at 600-630nm. Each butterfly sample was measured alongside a sample of female *Drosophila melanogaster* (Oregon-R strain, genome size of ≈ 175 Mb [79]) to establish a reference genome position. Single nuclei were identified by plotting area versus width for the DNA labelling with 5-50k positive nuclei recorded. For analysis, G0/1 peaks were gated for both the *D. melanogaster* and butterfly cells and relative intensities were then used to determine the genome size of the butterfly species using `FlowJo v9.6`.

### Life-history, karyotype and geographic range data

Current census sizes were estimated as the product of geographic range and density. All species in this study can be found in the region of Catalonia, Spain, where butterfly monitoring has been taking place since 1994 [80]. Density estimates were calculated as the mean number of individuals of each species seen per transect where that species is found, per year. The area range of each species was estimated from GBIF occurrence data (see Supplementary Data 6). The R package *rbgif* [81] was used to retrieve occurrence records — human observations with complete latitude and longitude information — for each species. Convex polygon areas (*km*^2^) were calculated using the function *eoo* in the R package *red* [82]. For species with large ranges, this was done separately for each land mass (to avoid including large bodies of water).

A list of larval host plants (LHP) for each species was compiled from [83] and HOST database [84].

Species were characterised as monophagous when LHPs were limited to one family or polyphagous when LHPs represented multiple families. Mean forewing length (across at least ten individuals per sex) reported in [85] was used as a proxy for adult body size. The mean between sexes was used for statistical analysis. Estimates of egg volume were retrieved from [86], haploid chromosome number from [85] and information on voltinism from [83]. Since the number of generations can vary within species, we only classified species as monovoltine if they had strictly one generation per year throughout their European range and polyvoltine if otherwise. In species with variable chromosome numbers, the mean was used for statistical analyses. All data can be found in Supplementary Data 3.

## Acknowledgments

We thank Jarrod Hadfield for help with MCMCglmm and discussions throughout and for comments on an earlier version of this manuscript, Lisa Cooper for excellent work in the wetlab and Carla and Oskar Lohse for their enthusiastic support in the field. We are indebted to Cecilia Corbella, Michael Jowers, Karl Wotton, Luis Valledor, Martim Melo, Jose Campos and Megan Wallace for help with field and lab logistics, Paul Jay for contributing samples and Richard Lewington from Collins Butterfly Guide for permission to reproduce illustrations. Permissions for field sampling were obtained from the Generalitat de Catalunya (SF/639), the Gobierno de Aragon (INAGA/500201/24/2018/0614 to Karl Wotton) and the Gobierno del Principado de Asturias (014252). This project was supported by an ERC starting grant (ModelGenomLand) and an Independent Research fellowship from the Natural Environmental Research Council (NERC) UK (NE/L011522/1) to KL. AM was supported by a summer studentship from the Institute of Evolutionary Biology at Edinburgh University, AH is supported by a Biotechnology and Biological Sciences Research Council David Phillips fellowship (BB/N020146/1) and RV is supported by project CGL2016-76322-P (AEI/FEDER, UE).

---

**Table S1.** Posterior mean estimates of the slope of linear correlates of genetic diversity at synonymous (*π*_4*D*_) and non-synonymous (*π*_0*D*_) sites inferred under a maximal model. For discrete predictors (LHP breadth and voltinism) the level of the factor described in the tabel is indicated in backets

| Predictor | Response | Posterior mean slope | 95% CI | $p_{MCMC}$ |
| --- | --- | --- | --- | --- |
| Current pop. size | $\pi_{4D}$ | 0.057 | -0.151, 0.263 | 0.602 |
| Current pop. size | $\pi_{0D}$ | 0.043 | -0.082, 0.182 | 0.530 |
| LHP breadth (poly.) | $\pi_{4D}$ | -0.303 | -0.860, 0.157 | 0.199 |
| LHP breadth (poly.) | $\pi_{0D}$ | -0.123 | -0.456, 0.198 | 0.412 |
| Body size | $\pi_{4D}$ | -0.269 | -0.524, -0.026 | 0.033 |
| Body size | $\pi_{0D}$ | -0.181 | -0.360, -0.023 | 0.030 |
| Relative egg size | $\pi_{4D}$ | -0.086 | -0.389, 0.195 | 0.551 |
| Relative egg size | $\pi_{0D}$ | 0.014 | -0.178, 0.201 | 0.873 |
| Voltinism (poly.) | $\pi_{4D}$ | 0.321 | -0.160, 0.775 | 0.151 |
| Voltinism (poly.) | $\pi_{0D}$ | 0.137 | -0.164, 0.441 | 0.348 |
| Chrom. number | $\pi_{4D}$ | 0.262 | 0.073, 0.471 | 0.012 |
| Chrom. number | $\pi_{0D}$ | 0.136 | 0.005, 0.268 | 0.044 |
| Genome size | $\pi_{4D}$ | 0.059 | -0.159, 0.302 | 0.609 |
| Genome size | $\pi_{0D}$ | 0.081 | -0.066, 0.231 | 0.250 |

**Fig S1.**
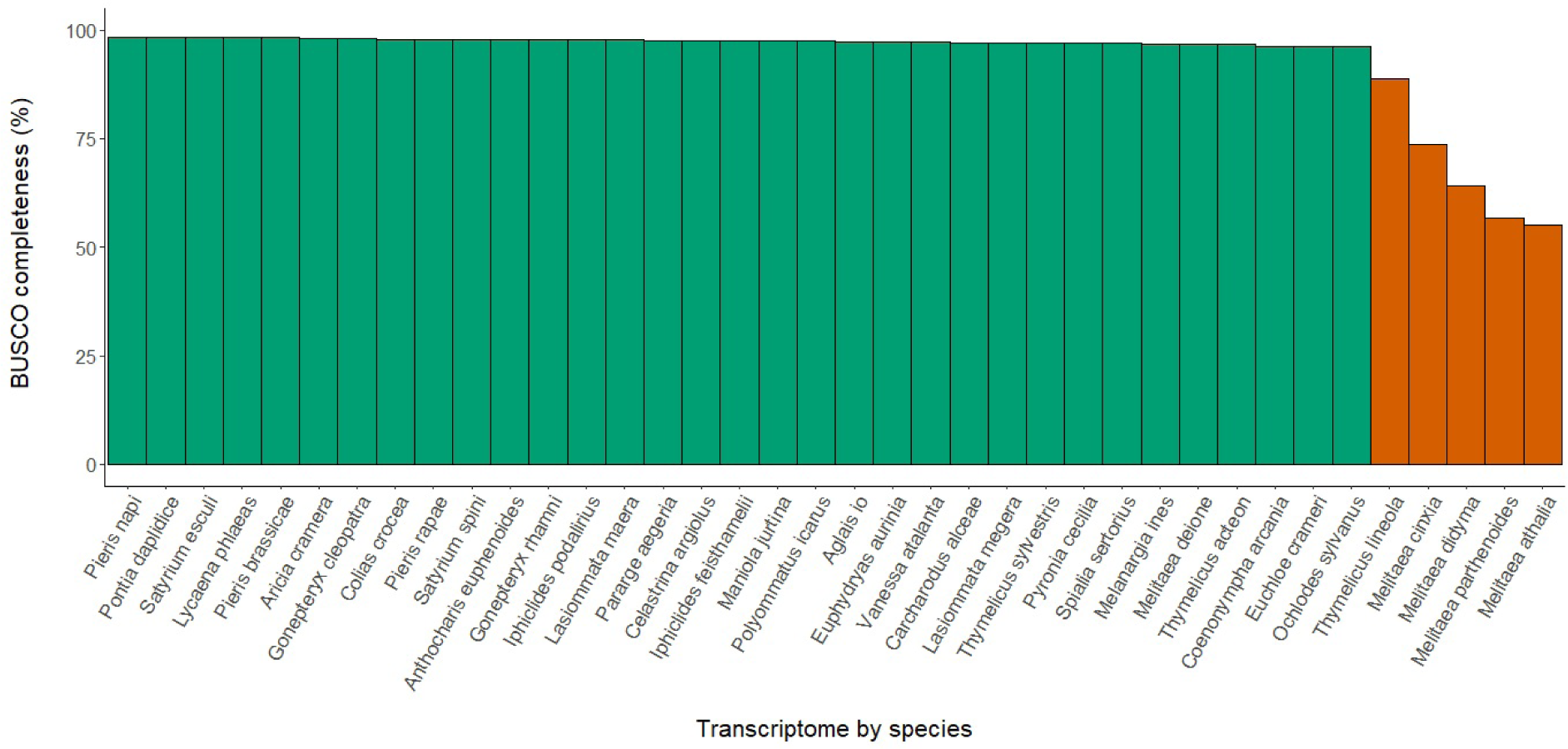
Completeness of transcriptomes assemblies were assessed using BUSCO scores. Transcriptomes assembled *de novo* as part of this study are shown in green, assemblies based on data from [5] in orange.

**Fig S2.**
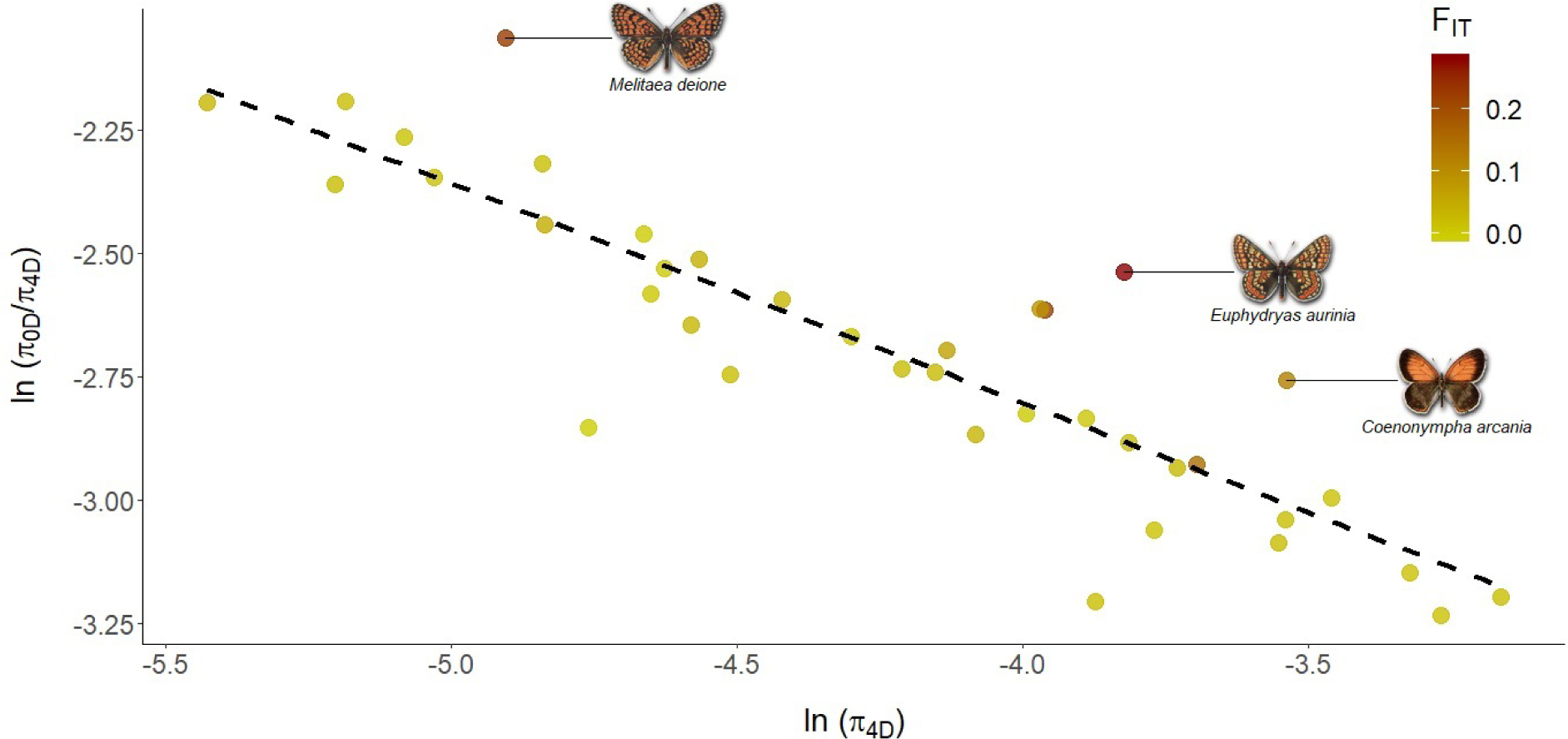
The log-log negative relationship between genetic diversity (*ln*(*π*_4*D*_)) and selection efficacy (*ln*(*π*_0*D*_/*π*_4*D*_)) is shown, where the slope of −0.44 corresponds to the *β* parameter of the DFE. Species with high genetic differentiation (*F_IT_*) fall above the line of best fit, i.e. they have less efficient selection than expected.

**Fig S3.**
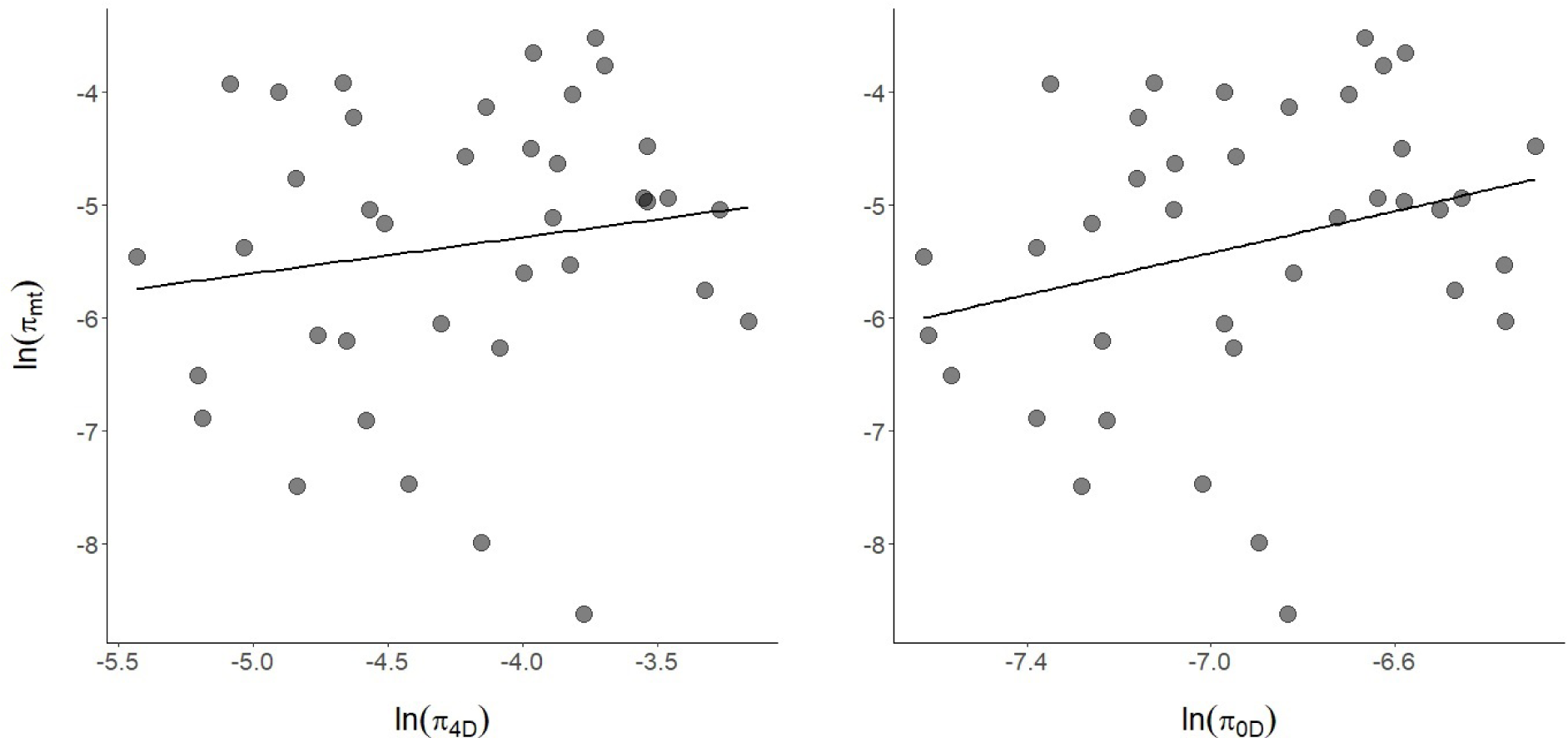
Mitochondrial diversity at the CO1 locus is essentially uncorrrelated with nuclear diversity both at synonymous (*π*_4*D*_, left) and non-synonymous (*π*_0*D*_, right) sites.

**Fig S4.**
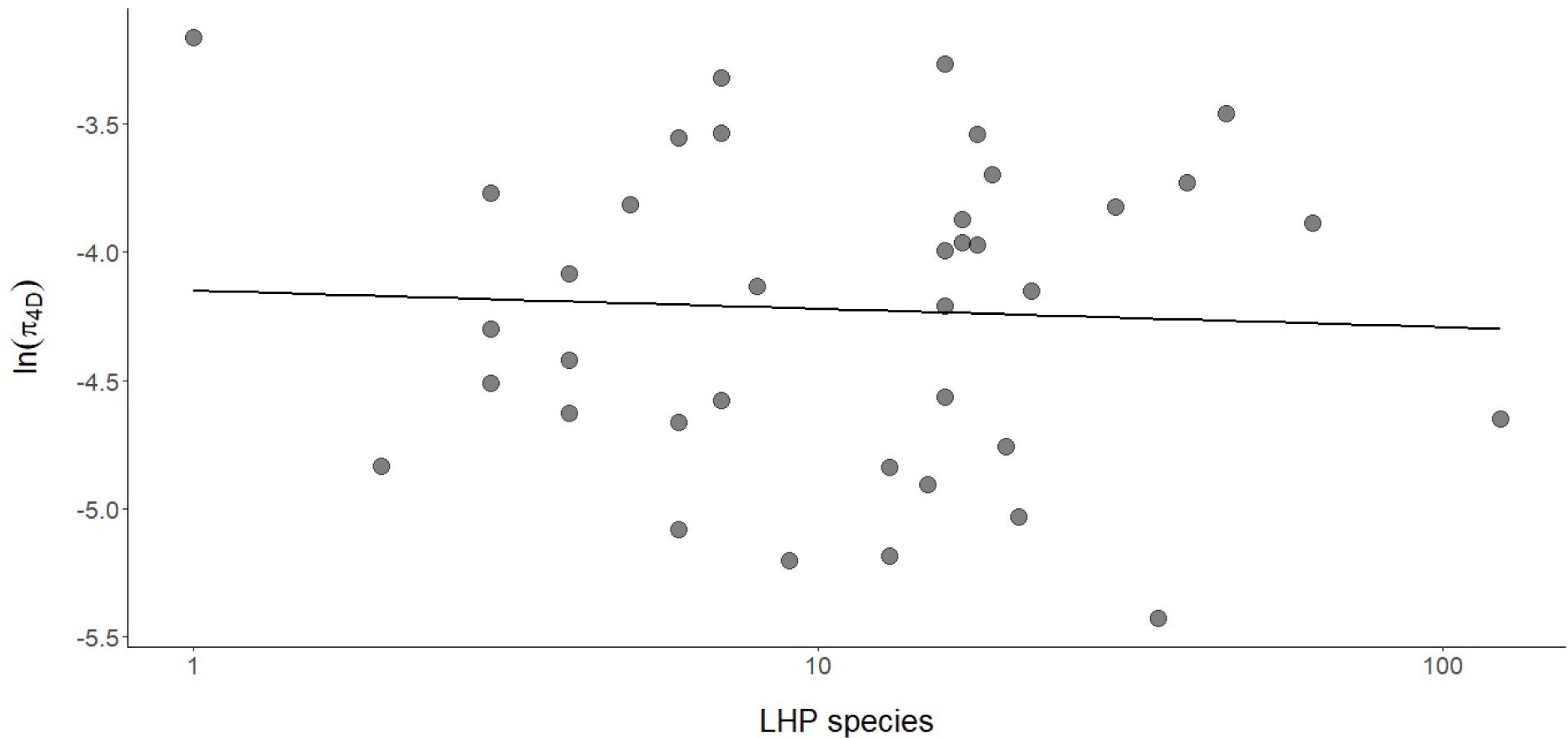
There is no correlation between the number of LHP species a butterfly species uses and its genome-wide genetic diversity.

**Fig S5.**
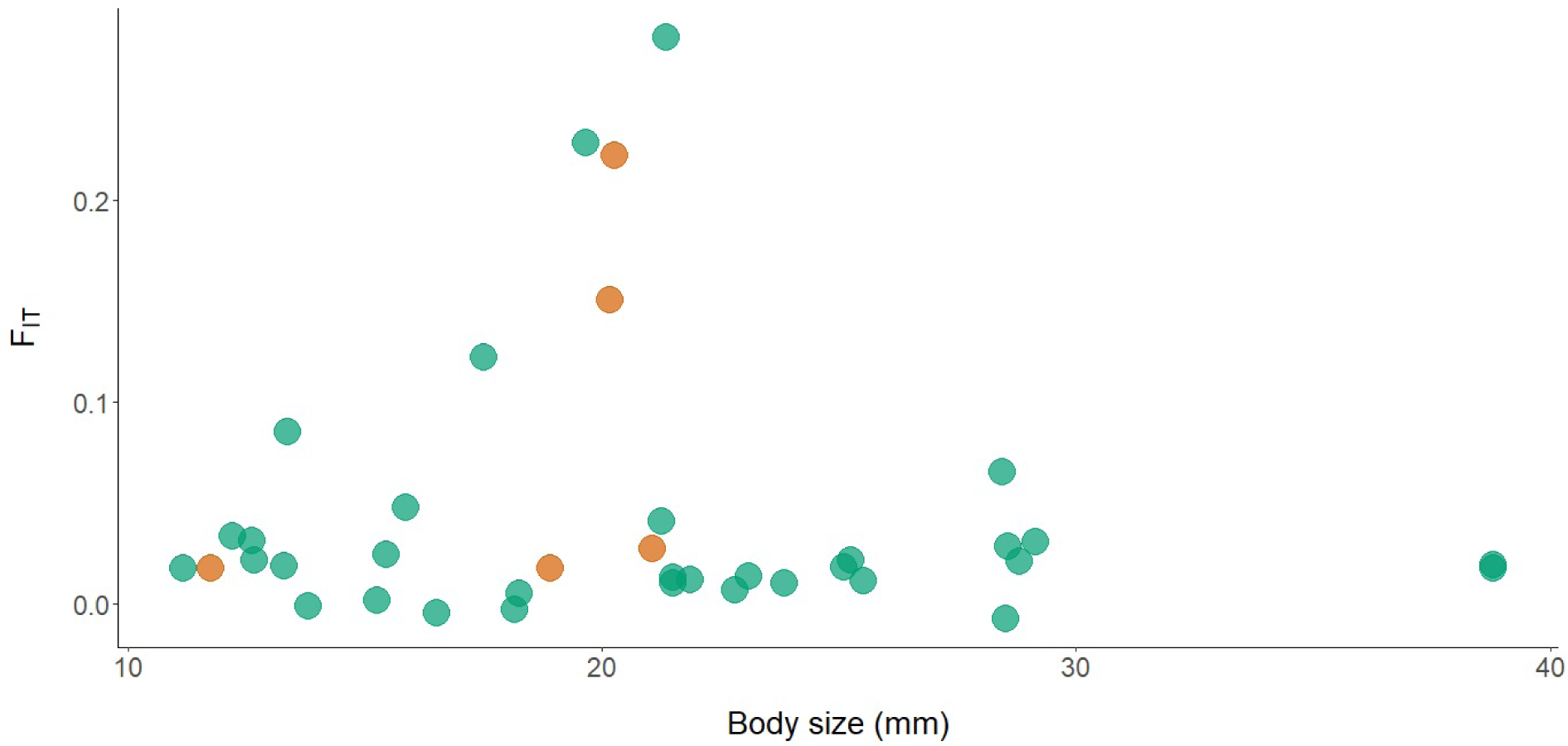
Genetic differentiation between individuals (*F_IT_*) sampled from different regions of Iberia is uncorrelated with body size. Species sampled within Iberia are shown in green and those sampled by [5] outside of Iberia are in orange.

**Fig S6.**
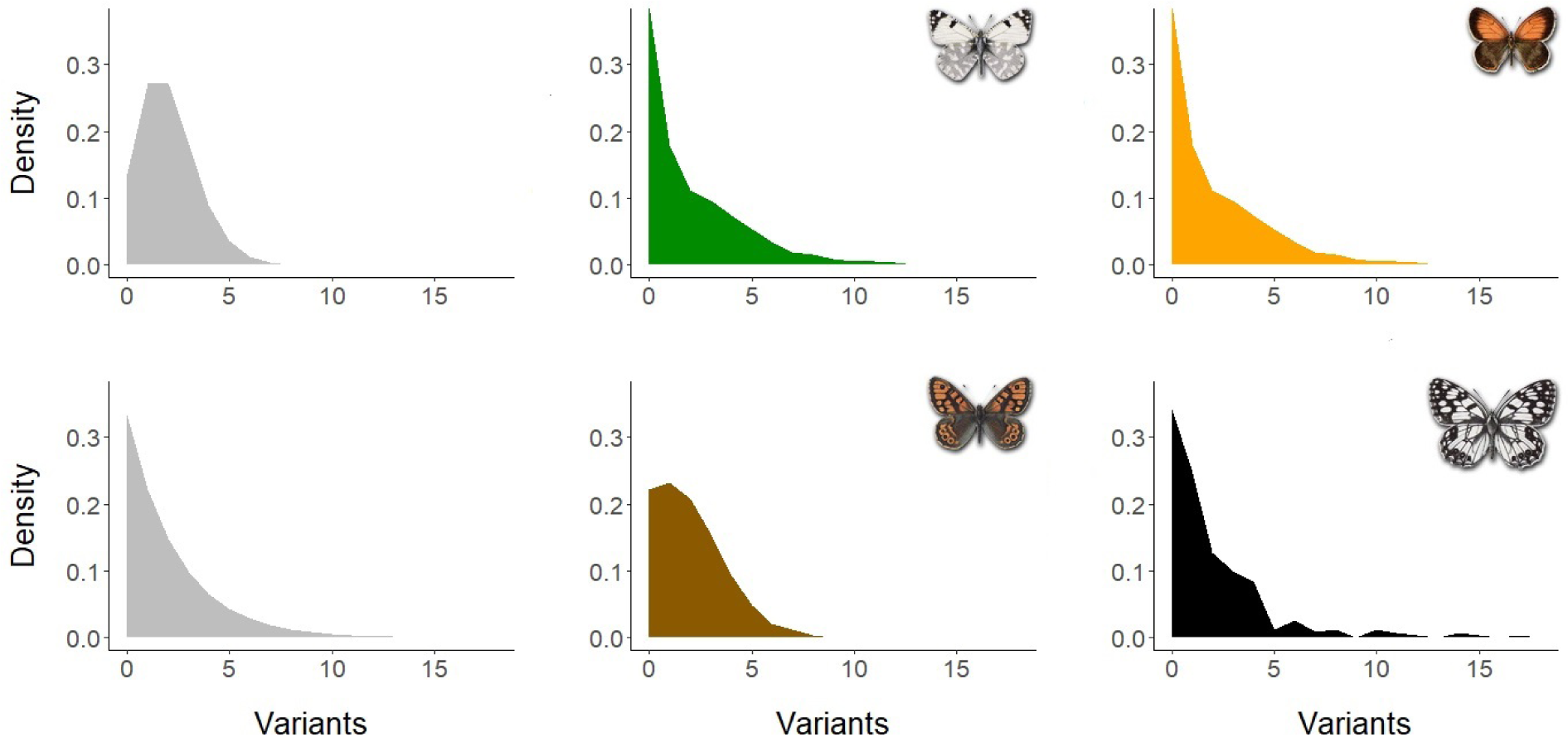
The distributions of heterozygous sites (*S*) in sequence blocks of a fixed length for different species: (top left) expectation under an extreme population expansion, (top middle) *Euchloe crameri* which is the median in the dataset, (top right) *Coenonympha arcania* the species closest to the expectation of constant *N_e_*, (bottom left) expectation under constant *N_e_*, (bottom middle) *Lasiommata megera* the species closest to the expectation of extreme expansion, (bottom right) *Melanargia ines* the species with the highest *V ar*[*S*] in the dataset.

**Fig S7.**
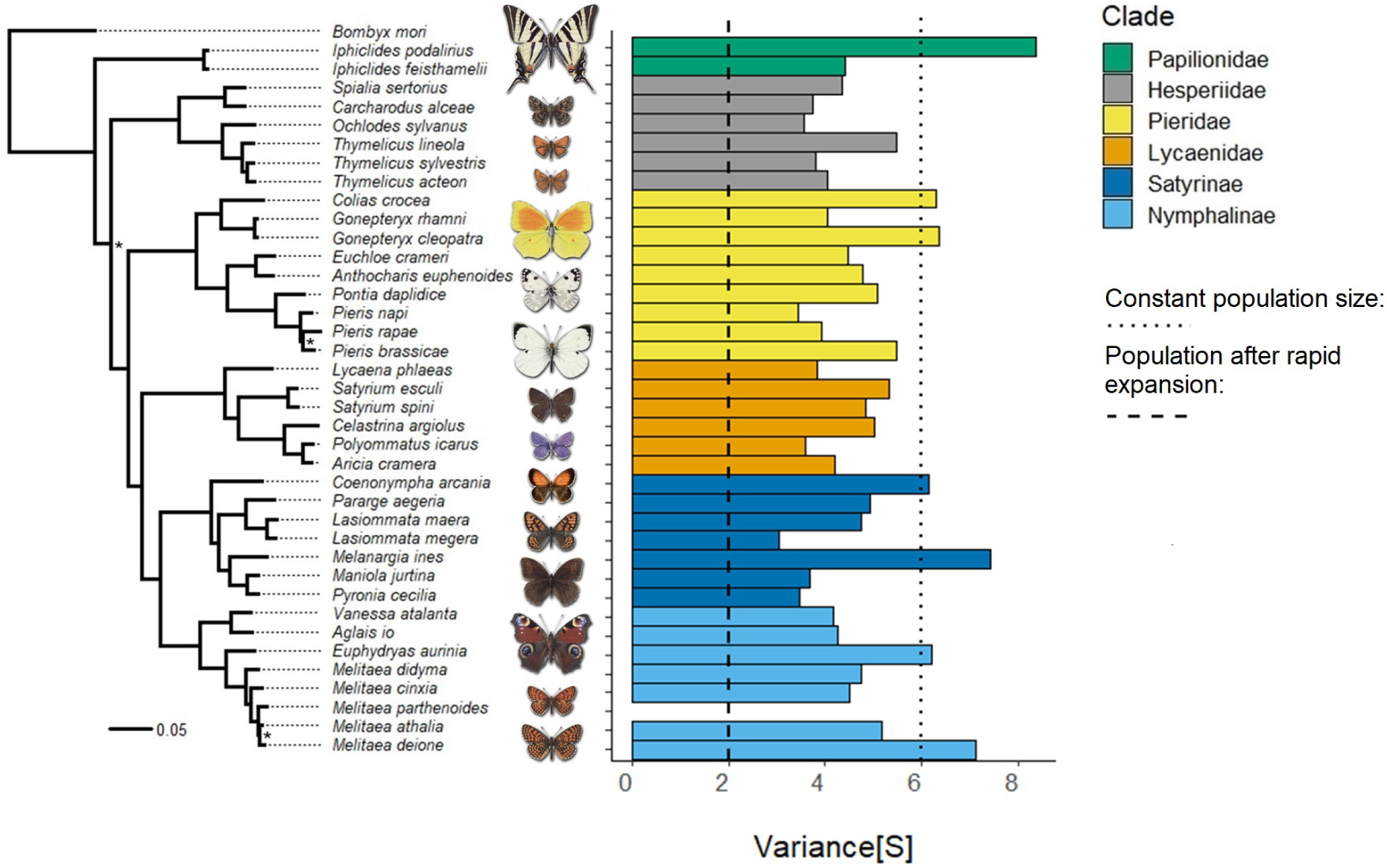
The variance in the number of heterozygous sites *V ar*[*S*] across the phylogeny. Vertical lines show the expected variance for a population of constant size and a rapidly expanding population. No estimate of *V ar*[*S*] is available for *Melitaea parthenoides* due to its low heterozygosity and transcriptome completeness.

**Fig S8.**
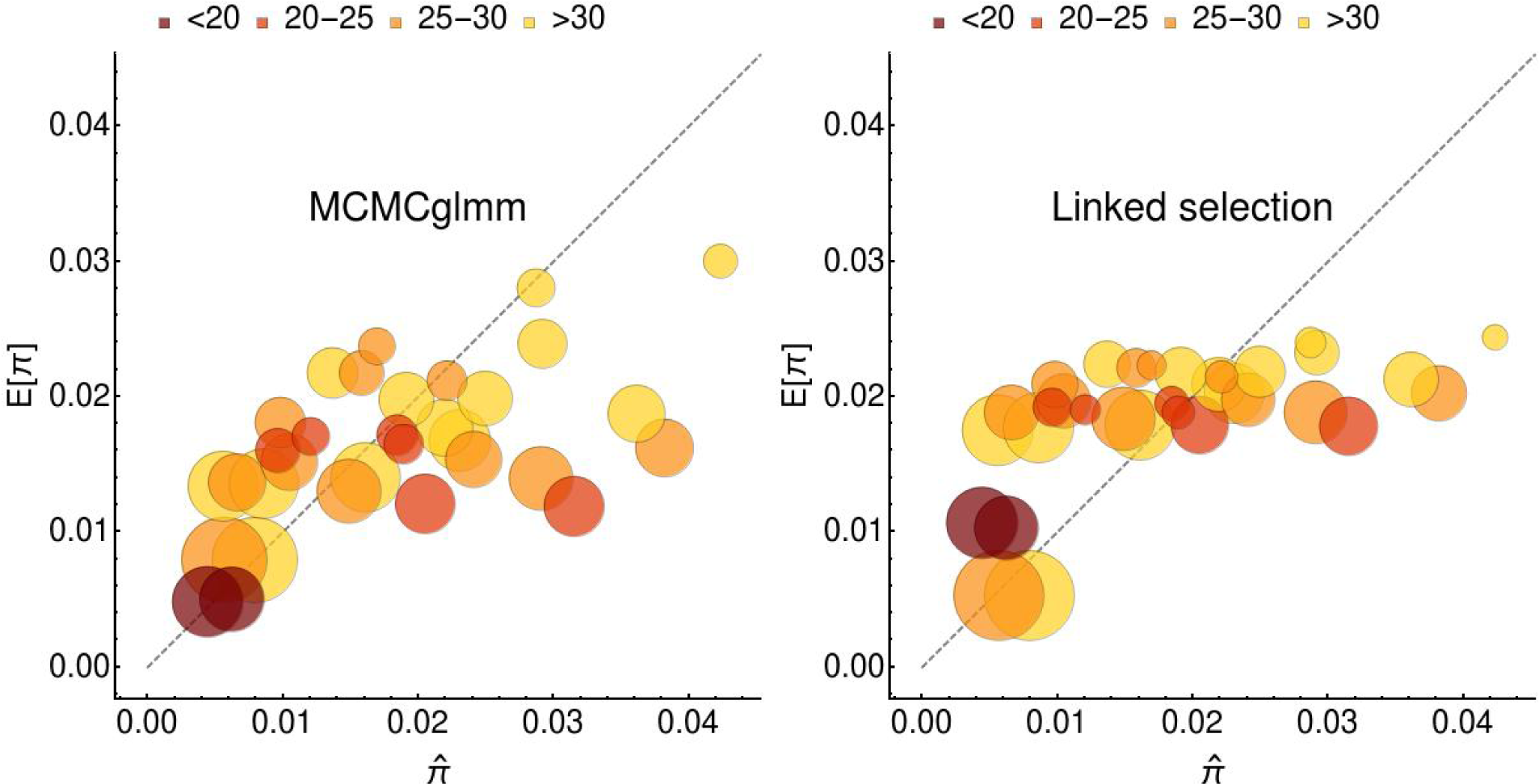
The minimal model inferred using MCMCglmm predicts the observed 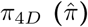 as well as an explicit model of effect of selection on linked neutral diversity. Circles are proportional to the relative body size of each species, the colour indicates chromosome number.

## References

1. Kimura M. The number of heterozygous nucleotide sites maintained in a finite population due to the steady flux of mutations. Genetics. 1969;61:893–903.

2. Watterson G. On the number of segregating sites in genetical models without recombination. TPB. 1975;7(2):256–276.

3. Kimura M. Theoretical foundation of population genetics at the molecular level. Theoretical Population Biology. 1971;2(2):174–208. Available from: http://www.sciencedirect.com/science/article/pii/0040580971900141.

4. Leffler EM, Bullaughey K, Matute DR, Meyer WK, Ségurel L, Venkat A, et al. Revisiting an old riddle: what determines genetic diversity levels within species? PLOS Biology. 2012 09;10(9):1–9. Available from: https://doi.org/10.1371/journal.pbio.1001388.

5. Romiguier J, Gayral P, Ballenghien M, Bernard A, Cahais V, Chenuil A, et al. Comparative population genomics in animals uncovers the determinants of genetic diversity. Nature. 2014;515:261–263.

6. Lewontin RC, Krakauer J. Distribution of gene frequency as a test of the theory of selective neutrality of polymorphisms. Genetics. 1973;74(1):175–195. Available from: http://www.genetics.org/content/74/1/175.

7. Nevo E. Genetic variation in natural populations: Patterns and theory. Theoretical Population Biology. 1978;13(1):121–177. Available from: http://www.sciencedirect.com/science/article/pii/0040580978900394.

8. Charlesworth B, Jain K. Purifying Selection, Drift, and Reversible Mutation with Arbitrarily High Mutation Rates. Genetics. 2014;198(4):1587–1602. Available from: http://www.genetics.org/content/198/4/1587.

9. Eanes WF. Analysis of Selection on Enzyme Polymorphisms. Annual Review of Ecology and Systematics. 1999;30(1):301–326. Available from: https://doi.org/10.1146/annurev.ecolsys.30.1.301.

10. Lynch M. Evolution of the mutation rate. Trends in Genetics. 2010;(6):345–352. Available from: https://doi.org/10.1016/j.tig.2010.05.003.

11. Maynard-Smith J, Haigh J. The hitch-hiking effect of a favourable gene. Genetics Research. 1974;23(5-6):23–35.

12. Ellegren H, Galtier N. Determinants of genetic diversity. Nature Reviews Genetics. 2016;17(6):422–. Available from: http://dx.doi.org/10.1038/nrg.2016.58.

13. Chen J, Glémin S, Lascoux M. Genetic Diversity and the Efficacy of Purifying Selection across Plant and Animal Species. Molecular Biology and Evolution. 2017;34(6):1417–1428. Available from: http://dx.doi.org/10.1093/molbev/msx088.

14. Charlesworth B, Morgan MT, Charlesworth D. The effect of deleterious mutations on neutral molecular variation. Genetics. 1993;134(4):1289–1303. Available from: http://www.genetics.org/content/134/4/1289.

15. Hudson RR, Kaplan NL. Deleterious background selection with recombination. Genetics. 1995;141(4):1605–1617. Available from: http://www.genetics.org/content/141/4/1605.

16. Corbett-Detig RB, Hartl DL, Sackton TB. Natural Selection Constrains Neutral Diversity across A Wide Range of Species. PLOS Biology. 2015 04;13(4):1–25. Available from: https://doi.org/10.1371/journal.pbio.1002112.

17. Coop G. Does linked selection explain the narrow range of genetic diversity across species? bioRxiv. 2016;Available from: https://www.biorxiv.org/content/early/2016/03/07/042598.

18. Espeland M, Breinholt J, Willmott KR, Warren AD, Vila R, Toussaint EFA, et al. A Comprehensive and Dated Phylogenomic Analysis of Butterflies. Current Biology. 2018;28(5):770–778. Available from: https://doi.org/10.1016/j.cub.2018.01.061.

19. Ehrlich AH, Ehrlich PR. Reproductive strategies in the butterflies: I. Mating frequency, plugging, and egg number. Journal of the Kansas Entomological Society. 1978;p. 666–697.

20. Davey JW, Chouteau M, Barker SL, Maroja L, Baxter SW, Simpson F, et al. Major Improvements to the Heliconius melpomene Genome Assembly Used to Confirm 10 Chromosome Fusion Events in 6 Million Years of Butterfly Evolution. G3: Genes, Genomes, Genetics. 2016;6(3):695–708. Available from: http://www.g3journal.org/content/6/3/695.

21. Zhan S, Huang J, Guo Q, Zhao Y, Li W, Miao X, et al. An integrated genetic linkage map for silkworms with three parental combinations and its application to the mapping of single genes and QTL. BMC Genomics. 2009 Aug;10(1):389. Available from: https://doi.org/10.1186/1471-2164-10-389.

22. Nei M. Genetic distance between populations. The American Naturalist. 1972;106 (949):283–292. Available from: https://doi.org/10.1086/282771.

23. Welch JJ, Eyre-Walker A, Waxman D. Divergence and Polymorphism Under the Nearly Neutral Theory of Molecular Evolution. Journal of Molecular Evolution. 2008;67(4):418–426. Available from: https://doi.org/10.1007/s00239-008-9146-9.

24. Keightley PD, Pinharanda A, Ness RW, Simpson F, Dasmahapatra KK, Mallet J, et al. Estimation of the Spontaneous Mutation Rate in Heliconius melpomene. Molecular Biology and Evolution. 2015;32(1):239–243. Available from: http://dx.doi.org/10.1093/molbev/msu302.

25. Kimura M. The Neutral Theory of Molecular Evolution. Cambridge University Press; 1983.

26. Ohta T. Slightly Deleterious Mutant Substitutions in Evolution. Nature. 1973;246(1):96–98. Available from: https://doi.org/10.1038/246096a0.

27. Campos JL, Zhao L, Charlesworth B. Estimating the parameters of background selection and selective sweeps in Drosophila in the presence of gene conversion. Proceedings of the National Academy of Sciences. 2017;114(24):E4762–E4771. Available from: https://www.pnas.org/content/114/24/E4762.

28. Loewe L, Charlesworth B. Inferring the distribution of mutational effects on fitness in *Drosophila*. Biology Letters. 2006;2(3):426–430.

29. Hewitt G. The genetic legacy of the Quaternary ice ages. Nature. 2000;405:907–913.

30. Schmitt T. Molecular biogeography of Europe: Pleistocene cycles and postglacial trends. Frontiers in Zoology. 2007 Apr;4(1):11. Available from: https://doi.org/10.1186/1742-9994-4-11.

31. E James J, Castellano D, Eyre-Walker A. DNA sequence diversity and the efficiency of natural selection in animal mitochondrial DNA. Heredity. 2016 11;118.

32. Nabholz B, Glémin S, Galtier N. The erratic mitochondrial clock: variations of mutation rate, not population size, affect mtDNA diversity across birds and mammals. BMC Evolutionary Biology. 2009 Mar;9(1):54. Available from: https://doi.org/10.1186/1471-2148-9-54.

33. Allio R, Donega S, Galtier N, Nabholz B. Large Variation in the Ratio of Mitochondrial to Nuclear Mutation Rate across Animals: Implications for Genetic Diversity and the Use of Mitochondrial DNA as a Molecular Marker. Molecular Biology and Evolution. 2017;34(11):2762–2772. Available from: http://dx.doi.org/10.1093/molbev/msx197.

34. Berlin Kolm S, Tomaras D, Charlesworth B. Low mitochondrial variability in birds may indicate Hill-Robertson effects on the W chromosome. Heredity. 2007 11;99:389–96.

35. Bazin E, Glémin S, Galtier N. Population Size Does Not Influence Mitochondrial Genetic Diversity in Animals. Science. 2006;312(5773):570–572. Available from: http://science.sciencemag.org/content/312/5773/570.

36. Barton NH. Genetic hitchhiking. Philosophical Transactions of the Royal Society of London B: Biological Sciences. 2000;355(1403):1553–1562. Available from: http://rstb.royalsocietypublishing.org/content/355/1403/1553.

37. Dincă V, Montagud S, Talavera G, Hernández-Roldán J, Munguira ML, García-Barros E, et al. DNA barcode reference library for Iberian butterflies enables a continental-scale preview of potential cryptic diversit. Scientific Reports. 2015;5:12395. Available from: https://www.nature.com/articles/srep12395.

38. Bunnefeld L, Hearn J, Stone GN, Lohse K. Whole-genome data reveal the complex history of a diverse ecological community. Proceedings of the National Academy of Sciences. 2018;115(28):E6507–E6515. Available from: http://www.pnas.org/content/115/28/E6507.

39. Talavera G, Lukhtanov VA, Rieppel L, Pierce NE, Vila R. In the shadow of phylogenetic uncertainty: the recent diversification of the Lysandra butterflies through chromosomal changes. Molecular Phylogenetics and Evolution. 2013;69:469–478.

40. Sjödin P, Kaj I, Krone S, Lascoux M, Nordborg M. On the Meaning and Existence of an Effective Population Size. Genetics. 2005;169(2):1061–1070. Available from: http://www.genetics.org/content/169/2/1061.

41. Bunnefeld L, Frantz LAF, Lohse K. Inferring bottlenecks from genome-wide samples of short sequence blocks. Genetics. 2015;201(3):1157–1169. Available from: http://www.genetics.org/content/201/3/1157.

42. White EP, Ernest SKM, Kerkhoff AJ, Enquist BJ. Relationships between body size and abundance in ecology. Trends in Ecology Evolution. 2007;22(6):323–330. Available from: http://www.sciencedirect.com/science/article/pii/S0169534707000985.

43. Brüniche-Olsen A, Kellner KF, Anderson CJ, DeWoody JA. Runs of homozygosity have utility in mammalian conservation and evolutionary studies. Conservation Genetics. 2018 Dec;19(6):1295–1307. Available from: https://doi.org/10.1007/s10592-018-1099-y.

44. Begun DJ, Aquadro CF. Levels of naturally occurring DNA polymorphism correlate with recombination rates in D. melanogaster. Nature. 1992;356(1):519–520. Available from: https://doi.org/10.1038.

45. Cutter DA, Payseur AB. Genomic signatures of selection at linked sites: Unifying the disparity among species. Nature reviews Genetics. 2013 03;14.

46. Ryan S, Lombaert E, Espeset A, Vila R, Talavera G, Dinca VE, et al. Global invasion history of the world’s most abundant pest butterfly: a citizen science population genomics study. 2018 12;.

47. Wiehe TH, Stephan W. Analysis of a genetic hitchhiking model, and its application to DNA polymorphism data from Drosophila melanogaster. Molecular Biology and Evolution. 1993;10(4):842–854. Available from: http://dx.doi.org/10.1093/oxfordjournals.molbev.a040046.

48. Coop G, Ralph P. Patterns of Neutral Diversity Under General Models of Selective Sweeps. Genetics. 2012;192(1):205–224. Available from: http://www.genetics.org/content/192/1/205.

49. Hill JA, Neethiraj R, Rastas P, Clark N, Morehouse N, de la Paz Celorio-Mancera M, et al. Cryptic, extensive and non-random chromosome reorganization revealed by a butterfly chromonome. bioRxiv. 2018;Available from: https://www.biorxiv.org/content/early/2018/03/02/233700.

50. Nash R, Brooker PC, Davis SJM. The Robertsonian translocation house-mouse populations of North East Scotland: A study of their origin and evolution. Heredity. 1983 06;50.

51. Patton JL, W Sherwood S. Chromosome Evolution and Speciation in Rodents. Annual Review of Ecology and Systematics. 2003 11;14:139–158.

52. Lewis H. The Origin of Diploid Neospecies in Clarkia. The American Naturalist. 1973;107(954):161–170. Available from: https://doi.org/10.1086/282824.

53. Šíchová J, Ohno M, Dincă V, Watanabe M, Sahara K, Marec F. Fissions, fusions, and translocations shaped the karyotype and multiple sex chromosome constitution of the northeast-Asian wood white butterfly, Leptidea amurensis. Biological Journal of the Linnean Society. 2016;118(3):457–471. Available from: http://dx.doi.org/10.1111/bij.12756.

54. Talla V, Suh A, Kalsoom F, Dincă V, Vila R, Friberg M, et al. Rapid Increase in Genome Size as a Consequence of Transposable Element Hyperactivity in Wood-White (Leptidea) Butterflies. Genome Biology and Evolution. 2017;9(10):2491–2505. Available from: http://dx.doi.org/10.1093/gbe/evx163.

55. Lynch M, Conery JS. The Origins of Genome Complexity. Science. 2003;302(5649):1401–1404. Available from: http://science.sciencemag.org/content/302/5649/1401.

56. Li J, Li H, Jakobsson M, Li S, Sjödin P, Lascoux M. Joint analysis of demography and selection in population genetics: where do we stand and where could we go? Molecular Ecology. 2012;21(1):28–44. Available from: https://onlinelibrary.wiley.com/doi/abs/10.1111/j.1365-294X.2011.05308.x.

57. Andrews S. FastQC a quality-control tool for high-throughput sequence data; 2015. Available from: http://www.bioinformatics.babraham.ac.uk/projects/fastqc/.

58. Ewels P, Magnusson M, Lundin S, Käller M. MultiQC: summarize analysis results for multiple tools and samples in a single report. Bioinformatics. 2016;32(19):3047–3048. Available from: http://dx.doi.org/10.1093/bioinformatics/btw354.

59. Bolger AM, Lohse M, Usadel B. Trimmomatic: a flexible trimmer for Illumina sequence data. Bioinformatics. 2014;30(15):2114–2120. Available from: http://dx.doi.org/10.1093/bioinformatics/btu170.

60. Haas BJ, Papanicolaou A, Yassour M, Grabherr M, Blood PD, Bowden J, et al. De novo transcript sequence reconstruction from RNA-seq using the Trinity platform for reference generation and analysis. Nature Protocols. 2013 Jul;8:1494 EP –. Available from: https://doi.org/10.1038/nprot.2013.084.

61. Simão FA, Waterhouse RM, Ioannidis P, Kriventseva EV, Zdobnov EM. BUSCO: assessing genome assembly and annotation completeness with single-copy orthologs. Bioinformatics. 2015;31(19):3210–3212. Available from: http://dx.doi.org/10.1093/bioinformatics/btv351.

62. Haas B, Papanicolaou A. Transdecoder (Find Coding Regions Within Transcripts);. Available from: https://github.com/TransDecoder/TransDecoder/wiki.

63. Altschul SF, Gish W, Miller W, Myers EW, Lipman DJ. Basic local alignment search tool. Journal of Molecular Biology. 1990;215(3):403–410. Available from: http://www.sciencedirect.com/science/article/pii/S0022283605803602.

64. Eddy SR, the HMMER development team. HMMER: biosequence analysis using profile hidden Markov models; 2018. Available from: http://hmmer.org/.

65. Li H. Aligning sequence reads, clone sequences and assembly contigs with BWA-MEM; 2013.

66. McKenna A, Hanna M, Banks E, Sivachenko A, Cibulskis K, Kernytsky A, et al. The Genome Analysis Toolkit: A MapReduce framework for analyzing next-generation DNA sequencing data. Genome Research. 2010;20(9):1297–1303. Available from: http://genome.cshlp.org/content/20/9/1297.abstract.

67. Quinlan AR, Hall IM. BEDTools: a flexible suite of utilities for comparing genomic features. Bioinformatics. 2010;26(6):841–842. Available from: http://dx.doi.org/10.1093/bioinformatics/btq033.

68. Garrison E, Marth G. Haplotype-based variant detection from short-read sequencing. ArXiv e-prints. 2012 Jul;.

69. Emms DM, Kelly S. OrthoFinder: solving fundamental biases in whole genome comparisons dramatically improves orthogroup inference accuracy. Genome Biology. 2015 Aug;16(1):157. Available from: https://doi.org/10.1186/s13059-015-0721-2.

70. Ratnasingham S, Hebert PDN. bold: The Barcode of Life Data System (http://www.barcodinglife.org). Mol Ecol Notes. 2007 May;7(3):355–364. PMC1890991[pmcid]. Available from: https://www.ncbi.nlm.nih.gov/pubmed/18784790.

71. Hall TA. BioEdit: a user-friendly biological sequence alignment editor and analysis program for Windows 95/98/NT. Nucleic Acids Symposium Series. 1999;41:95–98.

72. Thompson JD, Higgins DG, Gibson TJ. CLUSTAL W: improving the sensitivity of progressive multiple sequence alignment through sequence weighting, position-specific gap penalties and weight matrix choice. Nucleic Acids Res. 1994 Nov;22(22):4673–4680. 7984417[pmid]. Available from: https://www.ncbi.nlm.nih.gov/pubmed/7984417.

73. Stecher G, Kumar S, Tamura K. MEGA7: Molecular Evolutionary Genetics Analysis Version 7.0 for Bigger Datasets. Molecular Biology and Evolution. 2016 03;33(7):1870–1874. Available from: https://dx.doi.org/10.1093/molbev/msw054.

74. Katoh K, Standley DM. MAFFT Multiple Sequence Alignment Software Version 7: Improvements in Performance and Usability. Molecular Biology and Evolution. 2013;30(4):772–780. Available from: http://dx.doi.org/10.1093/molbev/mst010.

75. Capella-Gutiérrez S, Silla-Martínez JM, Gabaldón T. trimAl: a tool for automated alignment trimming in large-scale phylogenetic analyses. Bioinformatics (Oxford, England). 2009 August;25(15):1972—1973. Available from: http://europepmc.org/articles/PMC2712344.

76. Stamatakis A. RAxML version 8: a tool for phylogenetic analysis and post-analysis of large phylogenies. Bioinformatics. 2014;30(9):1312–1313. Available from: http://dx.doi.org/10.1093/bioinformatics/btu033.

77. Hadfield J. MCMC Methods for Multi-Response Generalized Linear Mixed Models: The MCMCglmm R Package. Journal of Statistical Software, Articles. 2010;33(2):1–22. Available from: https://www.jstatsoft.org/v033/i02.

78. DeSalle R, Gregory TR, Johnston JS. Preparation of Samples for Comparative Studies of Arthropod Chromosomes: Visualization, In Situ Hybridization, and Genome Size Estimation. In: Molecular Evolution: Producing the Biochemical Data. vol. 395 of Methods in Enzymology. Academic Press; 2005. p. 460–488. Available from: http://www.sciencedirect.com/science/article/pii/S0076687905950258.

79. Bennett MD, Leitch IJ, Price HJ, Johnston JS. Comparisons with Caenorhabditis (~100 Mb) and Drosophila (~175 Mb) Using Flow Cytometry Show Genome Size in Arabidopsis to be ~157 Mb and thus 25 Larger than the Arabidopsis Genome Initiative Estimate of ~125 Mb. Annals of Botany. 2003;91(5):547–557. Available from: http://dx.doi.org/10.1093/aob/mcg057.

80. catalanbms org; 2018. Available from: http://www.catalanbms.org.

81. Chamberlain S, Boettiger C. R Python, and Ruby clients for GBIF species occurrence data. PeerJ PrePrints. 2017;Available from: https://doi.org/10.7287/peerj.preprints.3304v1.

82. Cardoso P. red - an R package to facilitate species red list assessments according to the IUCN criteria. Biodiversity Data Journal. 2017;5:e20530. Available from: https://doi.org/10.3897/BDJ.5.e20530.

83. Tolman T, Lewington R. Collins Butterfly Guide. London: Harper Collins; 2009.

84. Robinson G, Ackery P, Kitching I, Beccaloni G, Hernandez L. HOSTS - A database of the the World’s Lepidopteran Hostplants; 2018. Available from: http://www.nhm.ac.uk/hosts.

85. Sanchez R, de los Angeles M. Volume 37: Lepidoptera: Papilionoidea. Fauna Iberica. London: Consejo Superior De Investigaciones Cientificas; 2013.

86. Garcia-Barros E. Body size, egg size, and their interspecific relationships with ecological and life history traits in butterflies (Lepidoptera: Papilionoidea, Hesperioidea). Biological Journal of the Linnean Society. 2000;70(2):251–284. Available from: http://dx.doi.org/10.1111/j.1095-8312.2000.tb00210.x.

