## Supplementary Data 4 for "The determinants of genetic diversity in butterflies – Lewontin’s paradox revisited"

We use equation derived by Barton (2000, eq. 1), modified for the case of semi-dominant favourable mutation with selection coefficient $s$ when homozygous. This gives the reduction in neutral diversity immediately following a sweep, measured relative to its initial value, as $\Delta p \approx (2N_e s)^{-(4r/s)}$, where $r$ is the frequency of recombination between the neutral and selected sites (for alternative derivations, see Coop and Ralph (2012); Weissman and Barton (2012); Elyashiv et al. (2016)). This can be written as $\Delta p \approx (2N_e s)^{-(2r/J)}$, where $J = s/[2\ln(2N_e s)]$. Following Weissman and Barton (2012), assume that $r$ is given by a linear function of physical distance $z$ between sites, $r = r_c z$. This is reasonable if recombination is due solely to crossing over, and sweep effects extend over relative small distances, so that double crossing over can be neglected. If, however, gene conversion also contributes to recombination, as is likely, the effect of a sweep is considerably reduced compared with this expression (Campos et al., 2017).

Following Kaplan et al. (1989) and Wiehe and Stephan (1993), $\Delta p$ can be equated to the probability that a sweep results in a coalescent event. Such an event is assumed to be instantaneous compared with coalescent events caused by genetic drift, whose rate is $1/(2N_0)$ in the absence of selective effects. If adaptive substitutions occur at a rate $\nu$ per basepair per generation, and there is no Hill-Robertson interference among them, the rate of sweep-induced coalescent events at a focal neutral site caused by all sweeps to the right and left of this site can be derived by treating the chromosome as a continuum, and integrating over all contributing sites. Following Weissman and Barton (2012), this approach gives the net rate of sweep-induced coalescent events per unit per unit coalescent time ($2N_0$ generations) as

$$C_s = 2N_0\nu \int_{-\infty}^{\infty} \exp(-2r_c z/J)\mathrm{d}z = 4N_0\nu \int_{0}^{\infty} \exp(-2r_c z/J)\mathrm{d}z = 2N_0\nu J r_c^{-1} \tag{1}$$

If we assume that there are $n_c$ chromosomes, each with a map length of 0.25 Morgans after taking the lack of crossing over in males into account, and a total haploid genome size of $G$ basepairs, we have $r_c = n_c/(4G)$. The total rate of substitutions per genome is $\nu_T = G\nu$. We thus have:

$$C_s \approx 8N_0 G\nu J n_c^{-1} = 8N_0 \nu_T J n_c^{-1} \tag{2}$$

### References


N. H. Barton. Genetic hitchhiking. *Philosophical Transactions of the Royal Society of London B: Biological Sciences*, 355(1403):1553–1562, 2000.

José Luis Campos, Lei Zhao, and Brian Charlesworth. Estimating the parameters of background selection and selective sweeps in Drosophila in the presence of gene conversion. *Proceedings of the National Academy of Sciences*, 114(24): E4762–E4771, 2017.

Graham Coop and Peter Ralph. Patterns of neutral diversity under general models of selective sweeps. *Genetics*, 192 (1):205–224, 2012.

Eyal Elyashiv, Shmuel Sattath, Tina T. Hu, Alon Strutsovsky, Graham McVicker, Peter Andolfatto, Graham Coop, and Guy Sella. A genomic map of the effects of linked selection in Drosophila. *PLOS Genetics*, 12(8):1–24, 08 2016.

N L Kaplan, R R Hudson, and C H Langley. The "hitchhiking effect" revisited. *Genetics*, 123(4):887–899, 1989.

Daniel B. Weissman and Nicholas H. Barton. Limits to the rate of adaptive substitution in sexual populations. *PLOS Genetics*, 8(6):1–18, 06 2012.

T H Wiehe and W Stephan. Analysis of a genetic hitchhiking model, and its application to dna polymorphism data from drosophila melanogaster. *Molecular Biology and Evolution*, 10(4):842–854, 1993.
