## Supplementary Data 5 for "The determinants of genetic diversity in butterflies – Lewontin’s paradox revisited"

### The expected slope of ln[$\pi$] on n

Equation 2:

$$E[\pi] = \frac{\pi_0}{8\, N_0\, J\, \nu_T\, n_c^{-1} + B^{-1}} \quad (1)$$

Thus the expectation for ln($\pi$) is:

$$E[\ln(\pi)] = -\ln\left(8\, N_0\, J\, \nu_T\, n_c^{-1} + B^{-1}\right) + \ln(\pi_0) \quad (2)$$

The expected slope on $n_c$ is:

$$\frac{d\ln(\pi)}{dn_c} = \frac{8\, J\, N_0\, \nu_T}{n_c^2\left(\frac{1}{B} + \frac{8\, J\, N_0\, \nu_T}{n_c}\right)} \quad (3)$$

In the limit of high $\nu_T$, this goes to $\frac{1}{n_c}$:

In[234]:=
$$\text{Limit}\left[\frac{8\, J\, N_0\, \nu_T}{n_c^2\left(\frac{1}{B} + \frac{8\, J\, N_0\, \nu_T}{n_c}\right)},\ \nu_T \to \text{Infinity}\right]$$

Out[234]=
$$\frac{1}{n_c}$$

Assuming $n = 25$ chromosomes, an upper bound for the slope is 0.04. Assuming no BGS (B=1) and a very high rate of sweeps, say $\nu_T\, N_0$= 500,000 and J= $10^5$ (which corresponds to a rate of sweep induced coalescence $S^{-1} = 0.8$), gives a slope of 0.0246:

In[255]:=
$$\left(\frac{8\, J\, N\nu}{n^2\left(\frac{1}{B} + \frac{8\, J\, N\nu}{n}\right)}\right) /.\ \{N\nu \to 500\,000,\ B \to 1,\ n \to 25,\ J \to 10^{-5}\} \ //\ N$$

Out[255]= 0.0246154

### Can we predict genetic diversity in butterflies?

#### Data import

Importing the real data (a csv file with columns, body size, chromN, Obs[$\pi$] and E[$\pi$]):

In[272]:=
```
butterflyDat =
  Import["/home/konrad/Dropbox/Manuscripts/Mackintosh_Butterfly_Comparative/
     Supplementary_Data/size_chromN_Obs[div]_E[div].csv", "Data"];
```

In[273]:=
```
(butterdat = (Flatten[#] & /@ Thread[{Range[38], Drop[butterflyDat, 1]}])) //
  TableForm
```

Out[273]//TableForm=

| | | | | | | | | |
|---|---|---|---|---|---|---|---|---|
| 1 | Aglais | io | 28.3 | 29.3 | 28.8 | 31 | 0.00549354 | 0.[illegible] |
| 2 | Anthocharis | euphenoides | 17.8 | 18.7 | 18.25 | 31 | 0.0135611 | 0.[illegible] |
| 3 | Aricia | cramera | 12.5 | 12.7 | 12.6 | 23 | 0.0120066 | 0.[illegible] |
| 4 | Carcharodus | alceae | 12.1 | 13.2 | 12.65 | 31 | 0.0286804 | 0.[illegible] |
| 5 | Celastrina | argiolus | 15.4 | 15.1 | 15.25 | 24 | 0.00954428 | 0.[illegible] |
| 6 | Coenonympha | arcania | 17.1 | 17.9 | 17.5 | 32 | 0.0291032 | 0.[illegible] |
| 7 | Colias | crocea | 23.3 | 24.4 | 23.85 | 31 | 0.023045 | 0.[illegible] |
| 8 | Euchloe | crameri | 20.8 | 22.2 | 21.5 | 31 | 0.0360898 | 0.[illegible] |
| 9 | Euphydryas | aurinia | 19.6 | 23.1 | 21.35 | 30 | 0.021892 | 0.[illegible] |
| 10 | Gonepteryx | cleopatra | 28 | 29.1 | 28.55 | n/a | 0.0102574 | n/[illegible] |
| 11 | Gonepteryx | rhamni | 28.2 | 28.7 | 28.45 | 31.5 | 0.0160305 | 0.[illegible] |
| 12 | Iphiclides | feisthamelii | 36.9 | 40.7 | 38.8 | 30 | 0.00790276 | 0.[illegible] |
| 13 | Iphiclides | podalirius | 36.9 | 40.7 | 38.8 | 30 | 0.00559308 | 0.[illegible] |
| 14 | Lasiommata | maera | 24.9 | 25.6 | 25.25 | 28 | 0.0148306 | 0.[illegible] |
| 15 | Lasiommata | megera | 21.2 | 22.5 | 21.85 | 29 | 0.0381182 | 0.[illegible] |
| 16 | Lycaena | phlaeas | 13.5 | 14.1 | 13.8 | 24 | 0.0184231 | 0.[illegible] |
| 17 | Maniola | jurtina | 24.6 | 25.6 | 25.1 | 29 | 0.029051 | 0.[illegible] |
| 18 | Melanargia | ines | 25.1 | 25.9 | 25.5 | 13 | 0.00620616 | 0.[illegible] |
| 19 | Melitaea | athalia | 19.7 | 20.6 | 20.15 | 31 | 0.02484 | 0.[illegible] |
| 20 | Melitaea | cinxia | 19.4 | 21.1 | 20.25 | 31 | 0.0190487 | 0.[illegible] |
| 21 | Melitaea | deione | 19.4 | 19.9 | 19.65 | n/a | 0.00740905 | n/[illegible] |
| 22 | Melitaea | didyma | 20.4 | 21.7 | 21.05 | 27.5 | 0.0240192 | 0.[illegible] |
| 23 | Melitaea | parthenoides | 17.8 | 20 | 18.9 | n/a | 0.0109648 | n/[illegible] |
| 24 | Ochlodes | sylvanus | 14.8 | 16.1 | 15.45 | 29 | 0.0157185 | 0.[illegible] |
| 25 | Pararge | aegeria | 20.8 | 21.7 | 21.25 | 27.5 | 0.0103858 | 0.[illegible] |
| 26 | Pieris | brassicae | 29.2 | 29.1 | 29.15 | 15 | 0.00439232 | 0.[illegible] |
| 27 | Pieris | napi | 23.6 | 22.6 | 23.1 | 25 | 0.0314815 | 0.[illegible] |
| 28 | Pieris | rapae | 23 | 22.6 | 22.8 | 25 | 0.0204783 | 0.[illegible] |
| 29 | Polyommatus | icarus | 13.8 | 12.9 | 13.35 | 23 | 0.0188731 | 0.[illegible] |
| 30 | Pontia | daplidice | 21.2 | 21.8 | 21.5 | 26 | 0.00652899 | 0.[illegible] |
| 31 | Pyronia | cecilia | 16.7 | 19.6 | 18.15 | 28 | 0.00978532 | 0.[illegible] |
| 32 | Satyrium | esculi | 15.5 | 16.2 | 15.85 | n/a | 0.0079324 | n/[illegible] |
| 33 | Satyrium | spini | 16.3 | 16.7 | 16.5 | n/a | 0.00943186 | n/[illegible] |
| 34 | Spialia | sertorius | 10.6 | 11.7 | 11.15 | 31 | 0.0423305 | 0.[illegible] |
| 35 | Thymelicus | acteon | 11.6 | 12.8 | 12.2 | 28 | 0.0168586 | 0.[illegible] |
| 36 | Thymelicus | lineola | 11.3 | 12.2 | 11.75 | 29 | 0.0208261 | n/[illegible] |
| 37 | Thymelicus | sylvestris | 12.8 | 13.8 | 13.3 | 27 | 0.0220458 | 0.[illegible] |
| 38 | Vanessa | atalanta | 27.9 | 29.1 | 28.5 | 31 | 0.00856634 | 0.[illegible] |

Removing species for which we  lack an estimate for chromosome number or genome size (*T.lineola* ):

In[274]:=
```
butterdatfltrd = DeleteCases[Delete[butterdat, {36}], {_, _, _, _, _, _, "n/a", _, _}];
```

Normalizing and z - transforming wing length (by the mean):

In[275]:=
```
busize = #[[-4]] & /@ butterdatfltrd;
normsize = (busize - (Mean[busize])) / StandardDeviation[busize];
sizeAndChrom = {normsize, #[[-3]] & /@ butterdatfltrd} // Thread;
```

In[277]:=
```
GraphicsRow[{ListPlot[{#[[-3]], #[[-2]]} & /@ butterdatfltrd,
    AxesStyle → {{Thick, Black}, {Thick, Black}}, AxesLabel → {"# Chroms", "π"}],
  ListPlot[{Exp[normsize], #[[-2]] & /@ butterdatfltrd} // Thread, AxesStyle →
    {{Thick, Black}, {Thick, Black}}, AxesLabel → {"Exp[size (mm)]", "π"}],
  ListPlot[{normsize, #[[-2]] & /@ butterdatfltrd} // Thread,
   AxesStyle → {{Thick, Black}, {Thick, Black}},
   AxesLabel → {"size (mm)", "π"}]}, ImageSize → 900]
```

Out[277]=
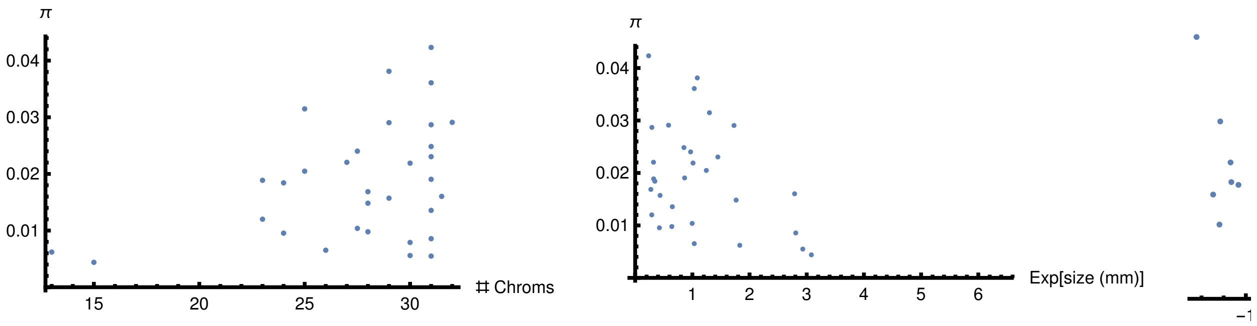


In[278]:=
```
πmcmcglmm = #[[-1]] & /@ butterdatfltrd; πreal = #[[-2]] & /@ butterdatfltrd;
```

### Checking the fit of a linear relationship between π and bodysize

Assuming that $N_e$ is a linear function of body size, the background $N_e$(across species) is 1.6 million!

In[279]:=
```
πlin[{nsize_, n_}] := 4 (Ne + nsize (β Ne)) μ;
```

In[280]:=
```
NeFittlin = Minimize[
  {Total[((πlin[#] & /@ sizeAndChrom) - πreal)^2] /. μ → 2.9 * 10^-9, {Ne > 0}}, {Ne, β}]
```

Out[280]=
```
{0.00269145, {Ne → 1.59193 × 10^6, β → -0.227077}}
```

The slope of -0.2 matches that found by MCMCglmm (as it should).

### BGS only & a linear model for $N_0$

We are assuming that $\pi_0$ is reduced by a factor $B = e^{-4\,U/n}$ where n is the number of chromosomes, i.e. the map length:

In[283]:=
```
πlinB[{nsize_, n_}] := 4 (Ne + nsize (β Ne)) μ * Exp[-((4 U) / n)];
```

Including BGS and assuming a maximum of U= 1 deleterious mutation per generation and genome only gives a slightly better fit.

In[284]:=
```
NeFittlinB = Minimize[
  {Total[((πlinB[#] & /@ sizeAndChrom) - πreal)^2] /. μ → 2.9 * 10^-9 /. U → 1, {Ne > 0}},
  {Ne, β}]
```

Out[284]=
```
{0.00255429, {Ne → 1.85953 × 10^6, β → -0.222689}}
```

In[285]:=
```
NeFittlinB[[1]] - NeFittlin[[1]]
```
Out[285]=
```
-0.000137157
```

U= 1 is an extremely conservative upper bound: assuming the Keightley et al. mutation rate for *Heliconius* (Keightley et al 2015) and a 400 Mb genome there are only 1.16 new mutations in total :
**Note**: Trying to estimate U without an upper bound gives a nonsensical result!

In[286]:=
```
NeFittlinBest =
 Minimize[{Total[((πlinB[#] &/@ sizeAndChrom) - πreal)^2] /. μ → 2.9 * 10^-9,
   {Ne > 0, U > 0}}, {Ne, β, U}]
```
Out[286]=
```
{0.00205356, {Ne → 1.10151 × 10^7, β → -0.257651, U → 13.4487}}
```

Assuming U=1, BGS would reduce diversity by a factor of B=0.74 to 0.88:

In[287]:=
```
N[Exp[-((4 U) / n)] /. {U -> 1, n → #}] &/@ {Min[#[[2]] &/@ sizeAndChrom],
  Mean[#[[2]] &/@ sizeAndChrom], Max[#[[2]] &/@ sizeAndChrom]}
```
Out[287]=
```
{0.735141, 0.865128, 0.882497}
```

### BGS AND Sweeps & a linear model for $N_0$

Fitting a model of BGS (with an input of U=1 deleterious mutations) and hard sweeps. The best fitting model assumes an unreasonably high rate of sweeps: $v$=0.134995 per generation!

In[288]:=
```
πlinBandSweeps[{nsize_, n_}] :=
  (4 (Ne + nsize (β Ne)) μ) / ((8 (Ne + nsize (β Ne)) * J (v / n)) + 1 / (Exp[-((4 U) / n)]));
```

In[289]:=
```
NeFittlinBandSweep = Minimize[
  {Simplify[Total[((πlinBandSweeps[#] &/@ sizeAndChrom) - πreal)^2] /. μ → 2.9 * 10^-9 /.
     U → 1 /. J → 10^-5], {Ne > NeFittlinB[[2, 1, 2]], 0.3 > v > 0}}, {Ne, β, v}]
```
Out[289]=
```
{0.00222163, {Ne → 5.88923 × 10^6, β → -0.356483, v → 0.135008}}
```

Ignoring BGS, so assuming hard sweeps only gives a slightly worse fit!

In[290]:=
```
NeFittlinSweep = Minimize[
  {Simplify[Total[((πlinBandSweeps[#] &/@ sizeAndChrom) - πreal)^2] /. μ → 2.9 * 10^-9 /.
     U → 0 /. J → 10^-5], {Ne > NeFittlinB[[2, 1, 2]], 0.2 > v > 0}}, {Ne, β, v}]
```
Out[290]=
```
{0.00223694, {Ne → 5.65451 × 10^6, β → -0.359324, v → 0.141397}}
```

If we measure model fit by the sum of least squares, the linked selection model fits (very slightly better) better than the linear model fitted by MCMCglmm and estimates a much higher background $N_e$!

In[291]:=
```
{Total[(πreal - πmcmcglmm)^2], NeFittlinBandSweep[[1]]}
```
Out[291]=
```
{0.00226291, 0.00222163}
```

However, if we plot the predicted vs observed $\pi$ reveals that the linked selection model predicts a much narrower range of genetic diversity than seen in the data:

In[292]:=
```
datlinSel =
  {πreal, (πlinBandSweeps[#] & /@ sizeAndChrom) /. μ → 2.9 * 10^-9 /. U → 1 /. J → 10^-5 /.
     NeFittlinBandSweep〚2〛, busize} // Thread;
datmcmcglmm = {πreal, πmcmcglmm, busize} // Thread;
```

In[293]:=
```
chrombin = {4, 4, 2, 4, 2, 4, 4, 4, 4, 4, 4,
   3, 3, 3, 2, 3, 1, 4, 4, 3, 3, 3, 1, 2, 2, 2, 3, 3, 4, 3, 3, 4};
```

In[294]:=
```
datlinSelBinned = Table[
   Drop[#, -1] & /@ SplitBy[SortBy[Flatten[#] & /@ ({datlinSel, chrombin} // Thread),
       #〚-1〛 &], #〚-1〛 &]〚i〛, {i, 1, 4}];
```

In[295]:=
```
datmcmcBinned = Table[
   Drop[#, -1] & /@ SplitBy[SortBy[Flatten[#] & /@ ({datmcmcglmm, chrombin} // Thread),
       #〚-1〛 &], #〚-1〛 &]〚i〛, {i, 1, 4}];
```

In[296]:=
```
datlinSelPlot = BubbleChart[datlinSelBinned, ChartBaseStyle → Opacity[0.7],
  Epilog → {Inset[Style["Linked selection", 30], {0.02, 0.034}]},
  FrameStyle → {{Thick, White}, {Thick, White}},
  FrameTicks → {{Automatic, None}, {Automatic, None}},
  FrameTicksStyle → Directive[Black], PlotRange → {{0, 0.043}, {0, 0.043}},
  FrameLabel → {"π̂", "E[π]"}, LabelStyle → Directive[Black, Large],
  BubbleSizes → {0.1, 0.35}, Prolog → {Thick, Dashed, Gray, Line[{{0, 0}, {1, 1}}]},
  ImageSize → 600, ChartStyle → "SolarColors",
  ChartLegends → Placed[{"<20", "20-25", "25-30", ">30"}, Top]]
datmcmcglmmPlot = BubbleChart[datmcmcBinned, ChartBaseStyle → Opacity[0.7],
  Epilog → {Inset[Style["MCMCglmm", 30], {0.02, 0.034}]},
  FrameStyle → {{Thick, White}, {Thick, White}},
  FrameTicks → {{Automatic, None}, {Automatic, None}},
  FrameTicksStyle → Directive[Black], PlotRange → {{0, 0.043}, {0, 0.043}},
  FrameLabel → {"π̂", "E[π]"}, LabelStyle → Directive[Black, Large],
  BubbleSizes → {0.1, 0.25}, Prolog → {Thick, Dashed, Gray, Line[{{0, 0}, {1, 1}}]},
  ImageSize → 600, ChartStyle → "SolarColors",
  ChartLegends → Placed[{"<20", "20-25", "25-30", ">30"}, Top]]
```

Out[296]=

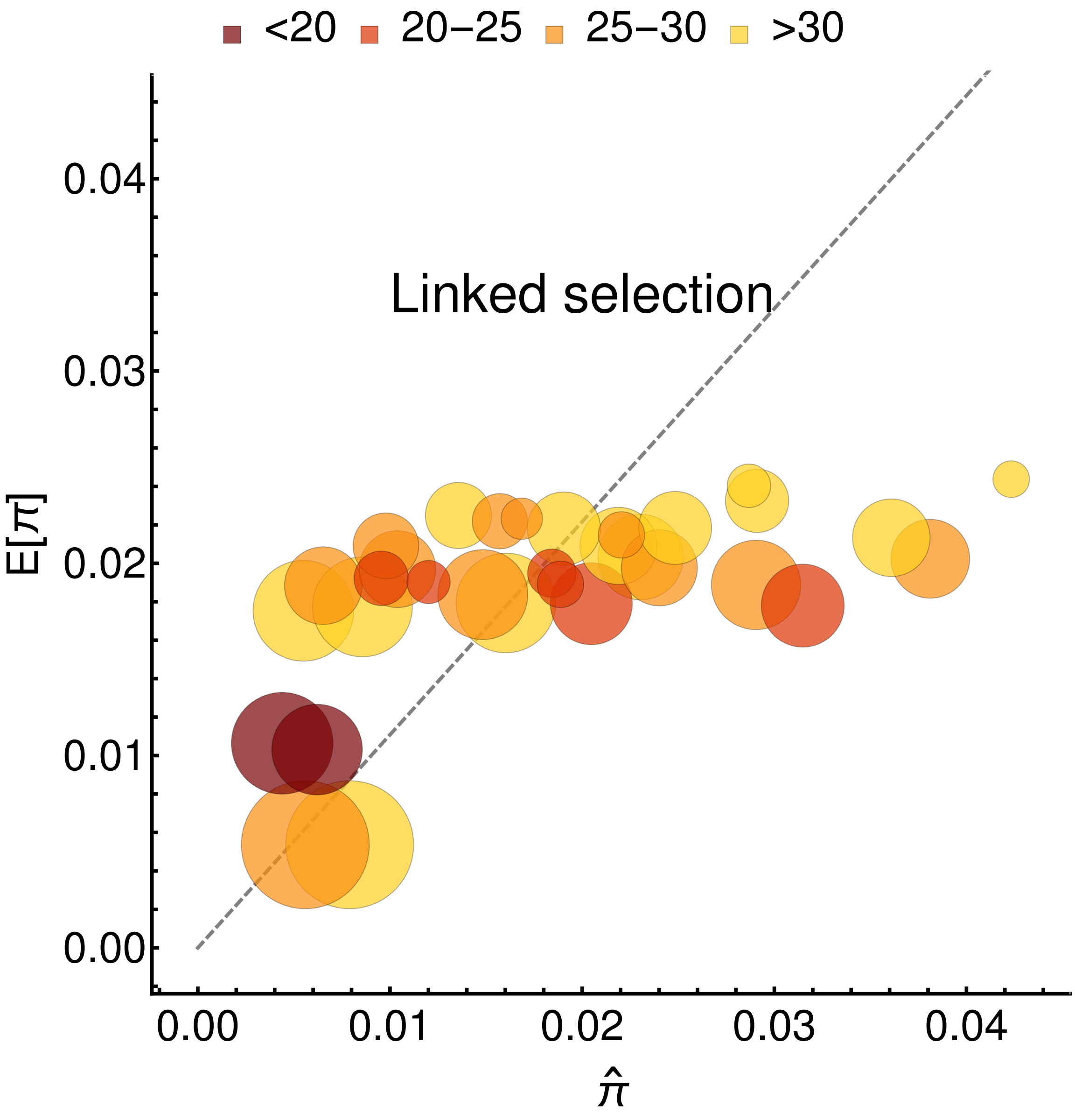

<20
20–25
25–30
>30
Linked selection
E[π]
0.04
0.03
0.02
0.01
0.00
0.00
0.01
0.02
0.03
0.04
π̂

Out[297]=

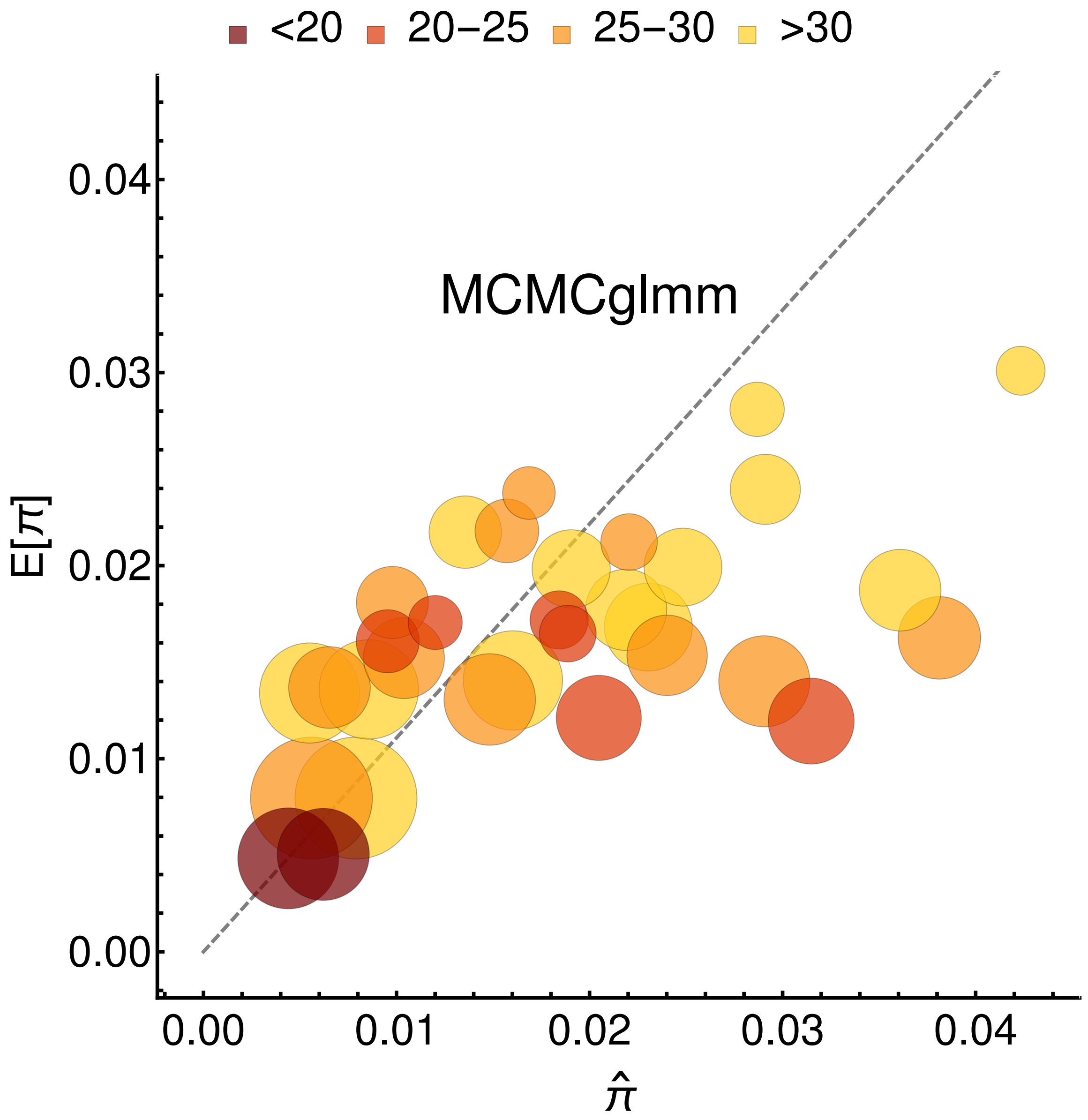


Circles are proportional to the body size, the color indicates chromosome number.

In[298]:=
```
Export[
  "/home/konrad/Dropbox/Manuscripts/Mackintosh_Butterfly_Comparative/figures/
    mcmcglmmPredict.jpg", datmcmcglmmPlot];
Export[
  "/home/konrad/Dropbox/Manuscripts/Mackintosh_Butterfly_Comparative/figures/
    datlinSelPlot.jpg", datlinSelPlot];
```

Given the assumed correlation between body size and $N_0$, genetic diversity depends only on the number of chromosomes for average (blue) or smaller (black) butterflies, but not the largest species in the set (green):

In[301]:=
```
Plot[{πlinBandSweeps[{Min[normsize], n}] /. NeFittlinBandSweep⟦2⟧ /. μ → 2.9 * 10^-9 /.
    U → 1 /. J → 10^-5,
  πlinBandSweeps[{0, n}] /. NeFittlinBandSweep⟦2⟧ /. μ → 2.9 * 10^-9 /. U → 1 /. J → 10^-5,
  πlinBandSweeps[{Max[normsize], n}] /. NeFittlinBandSweep⟦2⟧ /. μ → 2.9 * 10^-9 /.
    U → 1 /. J → 10^-5}, {n, 13, 32},
 PlotStyle → {Black, Blue, Green}, AxesLabel → {"chrom", "E[π]"},
 AxesStyle → {{Thick, Black, Large}, {Thick, Black, Large}},
 ImageSize → Large]
```

Out[301]=

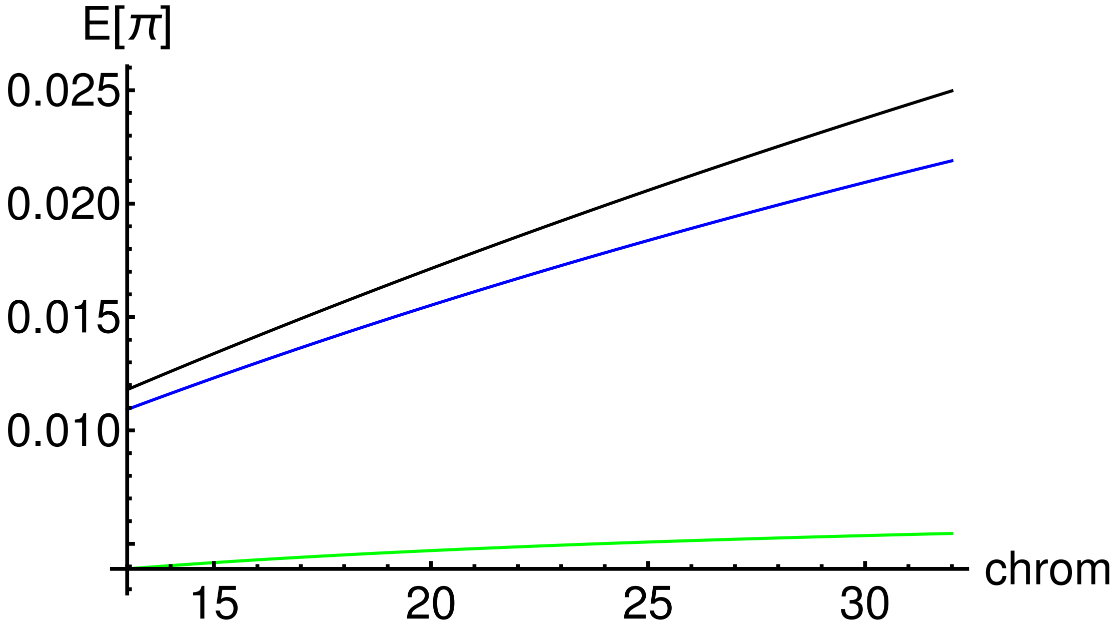
