## Supplementary Data 6 for "The determinants of genetic diversity in butterflies – Lewontin’s paradox revisited"

The following GBIF occurrence data were used to compute area ranges for 38 species of European butterfly :

### References


[1] de Vries H. Observation.org, Nature data from the Netherlands. Observation.org.; 2018. Available from: https://doi.org/10.15468/5nilie.

[2] Shah M CS. Artportalen (Swedish Species Observation System). Version 92.143. ArtDatabanken; 2019. Available from: https://doi.org/10.15468/kllkyl.

[3] T B. BioFokus. Version 1.966. Natural History Museum, University of Oslo.; 2019. Available from: https://doi.org/10.15468/jxbhqx.

[4] J R. Banco de Datos de la Biodiversidad de la Comunitat Valenciana. Biodiversity data bank of Generalitat Valenciana.; 2017. Available from: https://doi.org/10.15468/b4yqdy.

[5] M I. Programa de seguimiento de mariposas diurnas del PaÃ■s Vasco. Version 2.5. Basque Government.; 2018. Available from: https://doi.org/10.15470/7wmd2y.

[6] Caritg R MdlRS. ArtrÃšpodes d'Andorra. Version 1.3. Centre d'estudis de la neu i de la muntanya d'Andorra (CENMA), Institut d'Estudis Andorrans.; 2016. Available from: https://doi.org/10.15468/d9uwhe.

[7] Beja P CMGSJMFSSP Figueira R. EDP Foz Tua: Arthropoda âĂŞ Environmental Impact Assessment [2006-2008]. Version 1.6. EDP - Energias de Portugal.; 2018. Available from: https://doi.org/10.15468/jtdrhm.

[8] Vanreusel W VPGKSKDP Herremans M. Waarnemingen.be - Butterfly occurrences in Flanders and the Brussels Capital Region, Belgium. Version 1.6. Natuurpunt.; 2018. Available from: https://doi.org/10.15468/ezfbee.

[9] J B. The Distribution Atlas of Butterflies in Poland. Nicolaus Copernicus University of Torun.; 2017. Available from: https://doi.org/10.15468/yqzyas.

[10] F U. BioBlitz Barcelona 2010-14. Version 1.8. Museu de CiÃÍncies Naturals de Barcelona.; 2018. Available from: https://doi.org/10.15470/ssy7h3.

[11] Centre NBD. Butterflies of Ireland.; 2019. Available from: https://doi.org/10.15468/l7h1bv.

[12] Maes D BOVDHDP Brosens D. Vlinderdatabank - Butterflies in Flanders and the Brussels Capital Region, Belgium. Version 1.4. Research Institute for Nature and Forest (INBO).; 2017. Available from: https://doi.org/10.15468/njgbmh.

[13] Telenius A SM. Lepidoptera (Observations). GBIF-Sweden.; 2016. Available from: https://doi.org/10.15468/ao0ljg.

[14] I C. Atlas survey of the Butterflies of Denmark.; 2016. Available from: https://doi.org/10.15468/v5f2e2.

[15] C BE. Threatened species occurrences, Denmark 1991-2015. Version 1.4. Danish Nature Agency.; 2018. Available from: https://doi.org/10.15468/5cpovj.

[16] E MJ. Vegetation data from protected areas in Denmark (Âğ 3 in the Danish Nature Protection Act). Version 8.1. Department of Bioscience, Aarhus University.; 2016. Available from: https://doi.org/10.15468/ar7pbr.
